# Functional ultrasound imaging of stroke in awake rats

**DOI:** 10.1101/2023.05.31.543179

**Authors:** Clément Brunner, Gabriel Montaldo, Alan Urban

## Abstract

Anesthesia is a major confounding factor in preclinical stroke research as stroke rarely occurs in sedated patients. Moreover, anesthesia affects both brain functions and the stroke outcome acting as neurotoxic or protective agent. So far, no approaches were well suited to induce stroke while imaging hemodynamics along with simultaneous large-scale recording of brain functions in awake animals. For this reason, the first critical hours following the stroke insult and associated functional alteration remain poorly understood. Here, we present a strategy to investigate both stroke hemodynamics and stroke-induced functional alterations without the confounding effect of anesthesia, i.e., under awake condition. Functional ultrasound (fUS) imaging was used to continuously monitor variations in cerebral blood volume (CBV) in +65 brain regions/hemisphere for up to 3hrs after stroke onset. The focal cortical ischemia was induced using a chemo-thrombotic agent suited for permanent middle cerebral artery occlusion in awake rats, and followed by ipsi- and contralesional whiskers stimulation to investigate on the dynamic of the thalamo-cortical functions. Early (0-3hrs) and delayed (day 5) fUS recording enabled to characterize the features of the ischemia (location, CBV loss), spreading depolarizations (occurrence, amplitude) and functional alteration of the somatosensory thalamo-cortical circuits. Post-stroke thalamo-cortical functions were affected not only early after the stroke onset but were also altered secondarly and remotely from the initial insult. Overall, our procedure enables early, continuous, and chronic evaluations of hemodynamics and brain functions which, combined to stroke or other pathologies, aims to better understand physiopathologies toward the development of clinically relevant therapeutic strategies.

## Introduction

Stroke is a multifaceted and multiphasic pathology, complex to mimic under experimental conditions. Indeed, when compared to clinic, preclinical stroke models suffer from several limitations that narrow the experimental focus to few conditions^1–3^. Among these limitations, one can highlight the complexity to combine (i) imaging stroke in conscious animal models, (ii) addressing post-stroke brain functions and (iii) recording of hyperacute stroke hemodynamics, all crucial to design timely effective therapeutic strategies.

As first limitation, the use of anesthesia in preclinical studies seems to hamper the transition from animal to patient as most of stroke occurs in awake or sleeping patients^4,5^, but rarely in sedated patients. Moreover, anesthetics disrupt the brain functionality, alterates neurovascular coupling^6,7^, while differentially affecting metabolism, electrophysiology, temperature, blood pressure and tissue outcome by acting as neurotoxic or neuroprotective agents (see reviews^8–10^).

To date, only a few groups succeeded in inducing a stroke in awake rodents^11–14^. Moreover, post-stroke network and functional alterations have been addressed by few preclinical studies, providing evidence of functional network reorganization from minutes^15,16^ to days^17–20^ following stroke onset. However, these studies mostly focused on the cortical readouts and were unable to capture how deeper brain regions, like thalamic relays, were functionally and/or temporally affected remotely from the stroke insult (e.g., diaschisis)^21–23^. Furthermore, these studies were always conducted using various anesthetics (e.g., ventilated with halothane or isoflurane; medetomidine, urethane) known to differentially impact brain functions, as mentioned above.

Until recently, live monitoring of the hyperacute stroke-induced hemodynamics was restricted to few methods but often focused to the brain surface^13,24,25^. On the other hand, functional ultrasound (fUS), a recent neuroimaging modality measuring cerebral blood volume changes (CBV)^26–28^, was successfully employed to measure brain functions of awake rodents^29–34^, to address early post-stroke functional reorganization^16^, and to track stroke-induced hemodynamics at the brain-wide scale (i.e., ischemia and spreading depolarization)^35^. However, no study has further exploited such strategies to combine together stroke hemodynamics and brain-wide functional alteration in awake rodents.

In this study, we report on the stroke induction and the alteration of somatosensory brain functions in awake rats. Using the latest improvements toward imaging of awake rodents^29,31,33^ combined with chemo-thrombotic agent directly applied to the middle cerebral artery (MCA)^36,37^, we were able to induce MCA occlusion (MCAo) in awake rats while capturing continuous hemodynamic changes, including ischemia and spreading depolarization, in +65 brain regions for up to 3hrs after stroke onset. Finally, we investigated on how somatosensory thalamo-cortical functional reponses were progressively altered from early (0-3hrs) to late post-stroke (5d) timepoints.

## Materials and methods

### Animals

The experimental procedures were approved by the Committee on Animal Care of the Katholieke Universiteit Leuven, following the national guidelines on the use of laboratory animals and the European Union Directive for animal experiments (2010/63/EU). The manuscript was written according to the ARRIVE Essential 10 checklist for reporting animal experiments^38^. Adult male Sprague-Dawley rats weighed between 250–400g (n=9; Janvier Labs, France) were used. During habituation rats were housed two per cage kept in a 12-hr dark/light cycle at 23°C with *ad libitum* access to water and controlled access to food (15g/rat/day). After the initial surgical procedure, rats were housed alone. See **Supplementary Table 1** reporting on animal use, experimentation, inclusion/exclusion criteria.

### Body restraint and head fixation

The body restraint and head fixation procedures are adapted from published protocols and setup dedicated for brain imaging of awake rats^39–41^. Rats were habituated to the workbench and to be restrained in a sling suit (Lomir Biomedical inc, Canada) by progressively increasing restraining periods from minutes (5mins, 10mins, 30mins) to hours (1 and 3hrs) for one or two weeks. The habituation to head-fixation started by short (5 to 30s) and gentle head-fixation of the headpost between fingers. The headpost was then secured between clamps for fixation periods progressively increased following the same procedure as with the sling. For both body restraint and head fixation, the initial struggling and vocalization diminished over sessions. Water and food gel (DietGel, ClearH2O, USA) were provided for all body restraint and head-fixation habituation sessions. Once habituated, the cranial window for imaging was performed as described below (**Figure 1A-C**).

**Figure 1.**
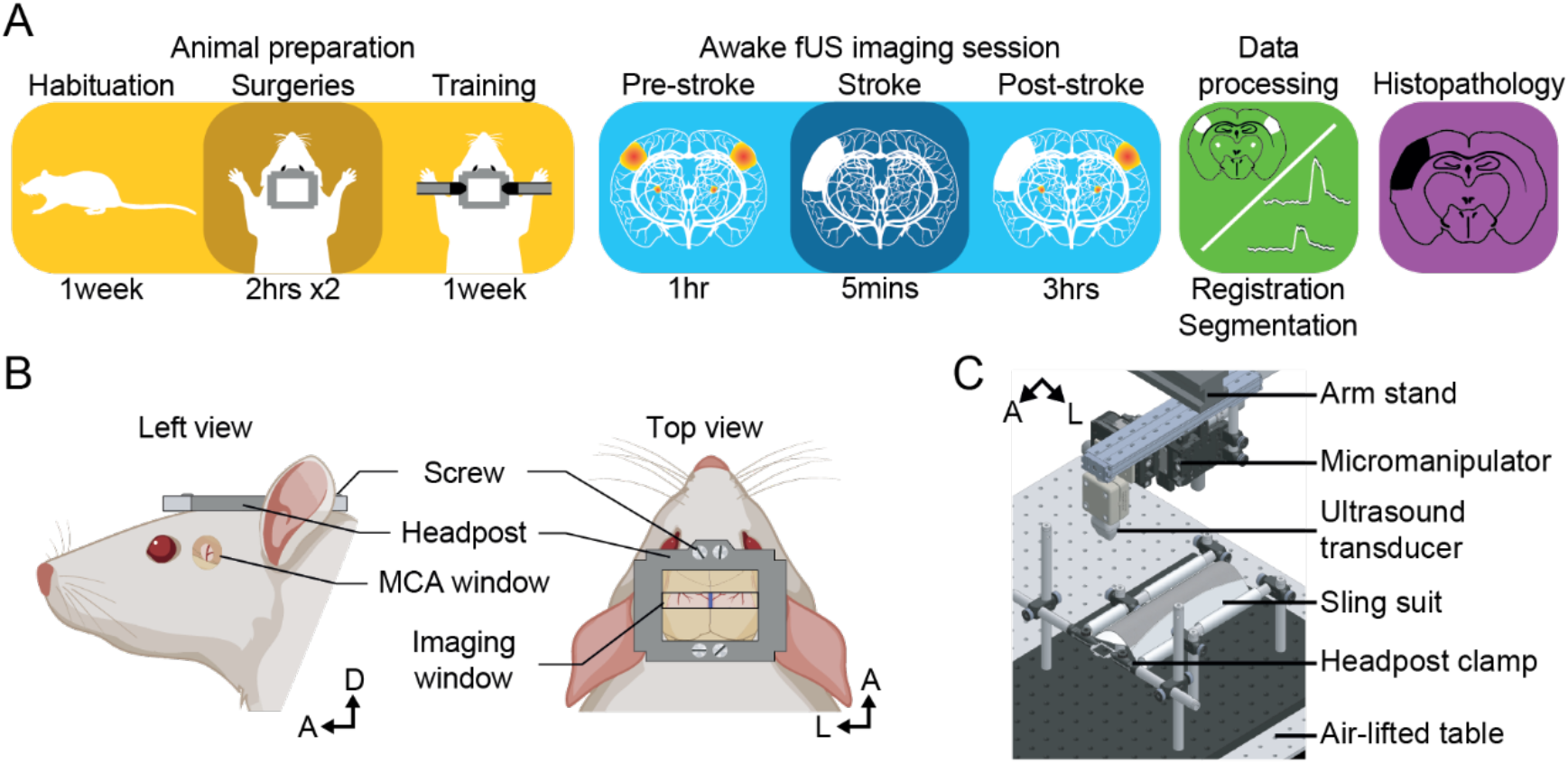
Experimental procedure. **(A)** Workflow for brain imaging of awake head-fixed rats including, from left to right: animal preparation (habituation to the bench, implantation of cranial windows, training), functional ultrasound (fUS) imaging of stroke induction and brain functions, data processing, and histopathology. **(B)** Overview of the headpost implantation and cranial windows developed for combined MCAo (left) and brain imaging (right) under awake conditions. **(C)** Computer-aided design of the experimental apparatus where the animal is placed and secured in a suspended sling suit and the head fixed by the means of clamps holding the headpost implanted to the rat skull. A, Anterior; D, Dorsal; L, Left.

### Surgical procedures

Cranial window over the MCA: Rats were anesthetized with isoflurane (5% for induction, 2% for maintenance; Iso-Vet, 1000 mg/g, Dechra, Belgium) and fixed in a stereotaxic frame. The depth of anesthesia was confirmed by the absence of reflex during paw pinching. After scalp removal and tissue cleaning, a 1-mm^2^ cranial window was performed at coordinates bregma +2mm and lateral 7mm, over the left distal branch of the MCA as reported in Brunner, Korostelev et al^16^. A silicone plug (Body Double-Fast Set, Smooth-on, Inc., USA) was used to protect the window and ease the access to the MCA before the occlusion procedure. Then, a stainless-steel custom designed headpost was fixed with bone screws (19010-00, FST, Germany) and dental cement (Super-Bond C&B, Sun Medical Co., Japan) to the animal skull (**Figure 1B**, left) as previously described by Brunner, Grillet et al.,^33^.

Cranial window for imaging: After recovery and habituation to head-fixation, a second cranial window was performed between bregma -2 to -4mm and 6mm apart from the sagittal suture (same anesthesia settings as the first cranial window; see above) following the procedure described in Brunner, Grillet et al.,^34^ (**Figure 1B**, right). This cranial window aims to cover bilateral thalamo-cortical circuits of the somatosensory whisker-to-barrel pathway. A silicone plug was also used to protect the window and a headshield was added to secure it^29^.

For both cranial windows, the dura mater was kept intact. After each surgery, rats were placed in their home cage and monitored until they woke up. Rats were medicated with analgesic (Buprenorphine, 0.1mg/kg, Ceva, France), anti-inflammatory (Dexamethasone, 0.5mg/kg, Dechra, Belgium) drugs injected directly after the surgery, at 24hrs and 48hrs after the surgery. An antibiotic (Emdotrim, 5%, Ecuphar, The Netherland) was added to the water bottle.

### Positionning

The mechanical fixation of the head-post ensures an easy and repeatabe positionning of the ultrasound probe across imaging session. The ultrasound probe is indeed fixed to a micromanipulator enabling light adjustements To find the plane of interest (containing both S1BF and thalamic relays: bregma - 3.4mm), we used brain landmarks (e.g., surface of the brain, hippocampus, superior sagittal sinus, large vessels). Note that as the headpost was carefully placed in the same position relative to the skulls landmarks (bregma and lambda), the position of the region of interest was minimal across animals

### Chemo-thrombotic stroke induction with ferric chloride solution

Once body restrained and head-fixed the silicone plug covering the MCA window was removed allowing the application of a drop of 20% ferric chloride solution (FeCl_3_; Sigma Aldrich, USA) to the MCA^36,37^ (**Figure 2**). Once the ischemia was visually detected using the real-time display of μDoppler images, the solution was washed out with saline to stop the reaction.

**Figure 2.**
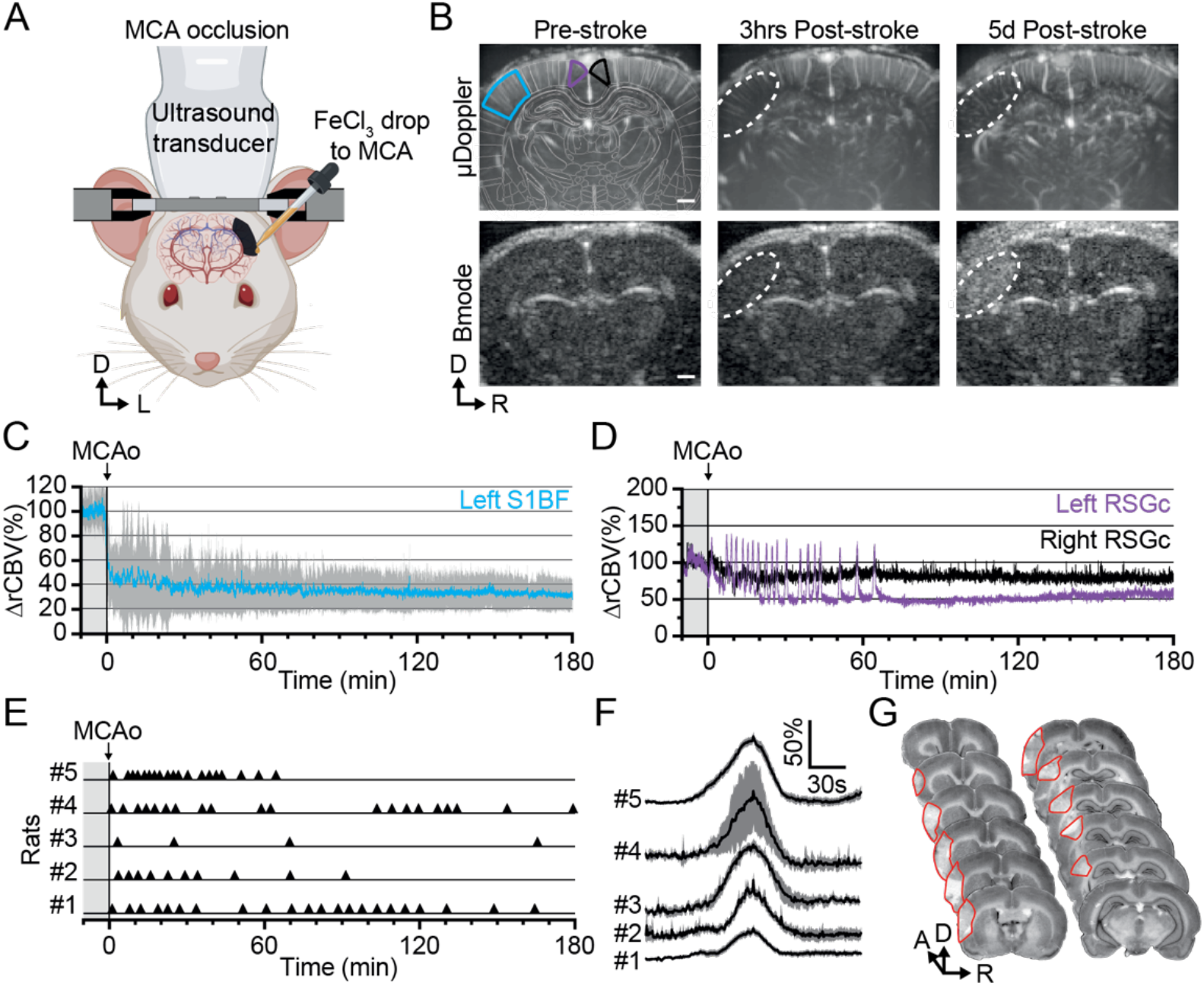
FeCl_3_-stroke induction under awake conditions. **(A)** Front view representation of functional ultrasound (fUS) imaging during live chemo-thrombosis of the left MCA with FeCl_3_ in awake head-fixed rats. **(B)** Set of typical coronal μDoppler images of the brain microvasculature (top row) and morphological Bmode images (bottom row) before stroke (left), 3hrs (middle), and 5d after stroke onset (right) from the same animal. μDoppler images (top left) were registered and segmented based on a digital version of the rat brain atlas (white outlines). Colored outlines (cyan, purple, and black) delineate regions of interest plotted in (C) and (D). The white dotted region of interest highlights the ischemia in μDoppler images (Top row) and tissue hyper-echogenicity in Bmode (Bottom row). **(C)** Temporal plot of the average signal (ΔrCBV (%), mean±95%CI, n=5) in the barrel-field primary somatosensory cortex (S1BF, cyan) from the left hemisphere, affected by the MCAo. **(D)** Temporal plots of the average signal (ΔrCBV (%)) in the retrosplenial granular cortex (RSGc) from the affected (purple) and non-affected hemisphere (black) from the same animal. **(E)** Occurrence of spreading depolarizations after MCAo. Each horizontal line represents one rat; each triangle marker depicts the occurrence of one spreading depolarization. **(F)** Temporal plots of the average signal change (ΔrCBV (%), mean±95%CI, respectively black line and gray band) of hemodynamic events associated with spreading depolarizations (centered on the peak) for each rat (#1 to 5). **(G)** Typical rat brain cross-sections stained by cresyl violet to evaluate the tissue infarction at 24hrs after FeCl_3_-induction occlusion of MCA. The infarcted territory is delineated in red. Scale bars: 1mm. D: Dorsal; L: left; R: right.

### Whisker stimulation paradigm

Two stimulation combs individually controlled by a stepper motor (RS Components, UK) were used to deliver mechanical 5-Hz sinusoidal deflection of ∼20° of amplitude for 5s, alternatively to left and right whisker pads. For each whisker pad, trials were spaced by a period of 1min and 20s without stimulation. Thus, the effective delay between two stimulations delivered to the same whisker pad is 80 seconds from start to start. The blocks of stimulation were continuously delivered throughout the imaging sessions, time-locked with the fUS acquisition (**Figure 3**) to allow the subsequent analysis of hemodynamic responses within the fUS time-series.

**Figure 3.**
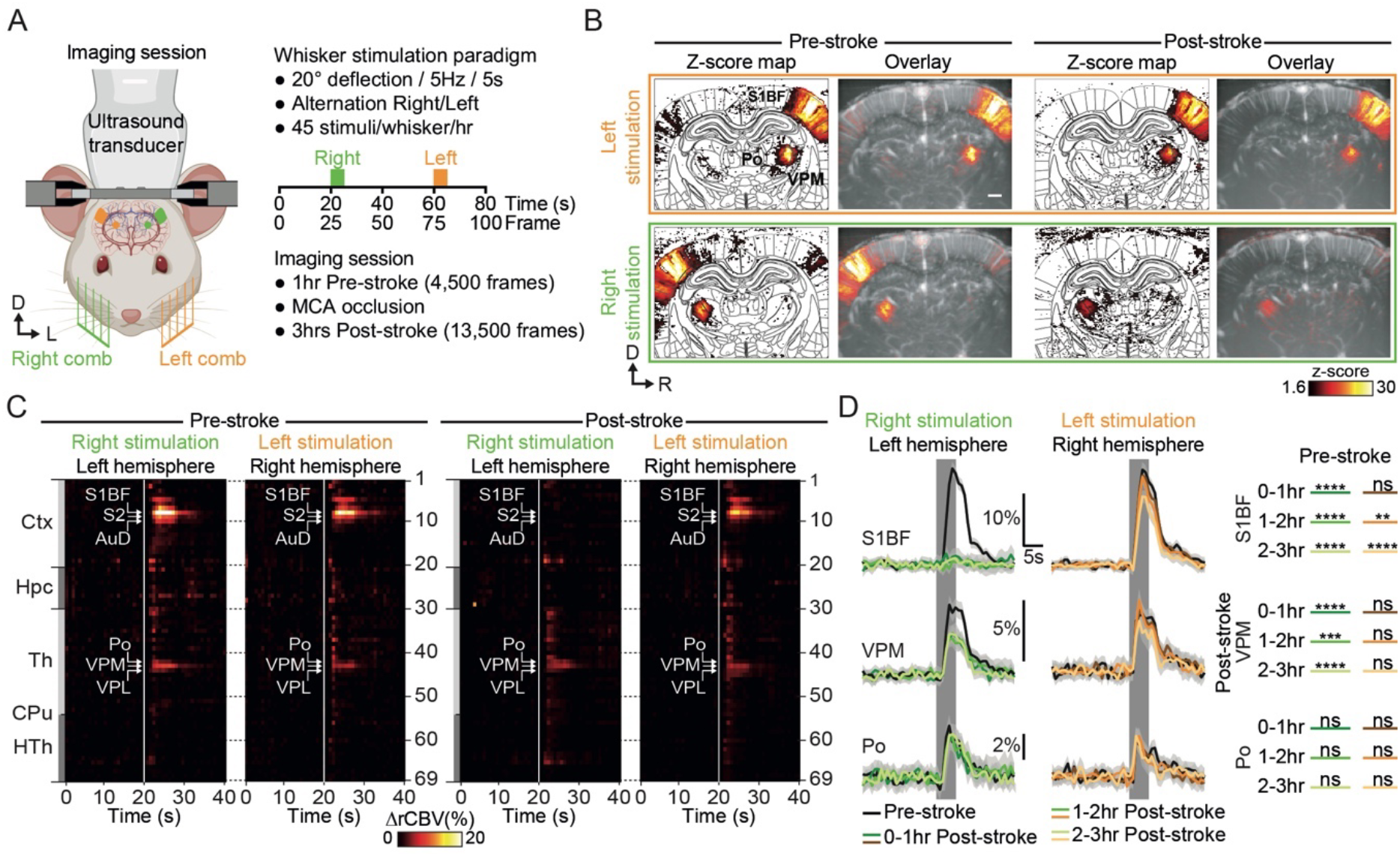
Early post-stroke alteration of whisker-to-barrel thalamo-cortical circuit. **(A**) Front view representation of functional ultrasound (fUS) imaging during repetitive stimulation of the left (orange) or right whisker pad (green) with a mechanical comb in awake head-fixed rats. Whisker stimulations were delivered alternately between left and right whisker pads before and early after MCAo. Each rat receives 45 stimuli per whisker pad each hour of imaging. **(B)** Average activity maps (z-score) from one rat depicting evoked functional responses to either left (orange) or right whisker pads stimulation (green) registered with a digital version of the rat Paxinos atlas (white outlines) and overlaid with the corresponding coronal μDoppler image, before (left; Pre-stroke, average of 45 trials) and after stroke induction in the left hemisphere (right; Post-stroke, average of 125 trials). **(C)** Region-time traces of the average hemodynamic changes (ΔrCBV(%)) in response to right (green) or left whisker stimulation (orange) extracted from the contralateral hemisphere (left and right, respectively) before (left; Pre-stroke, n=5, 45 trials/rat) and after stroke induction in the left hemisphere (right; Post-stroke, n=5, 135 trials/rat). Brain regions are ordered by major anatomical structures (see **Supplementary Table 2**). The vertical line represents the stimulus start. S1BF, S2, AuD, VPM, VPL and Po regions are brain regions significatively activated (all pvalue<0.01; GLM followed by t-test). A larger version of panel C is provided in **Supplementary Figure 3. (D)** Left, Average response curves from the S1BF, the VPM, and Po regions before (Pre-stroke, black, n=5, 45 trials/rat), and from first to third hour after stroke induction (0-1hr, 1-2hr, 2-3hr Post-stroke, orange and green, n=5, 45 trials/hr/rat). Data are mean±95%CI. The vertical bar represents the whisker stimulus. Right, Statistical comparison of the AUC between pre-stroke and post-stroke response curves for S1BF, VPM, and Po regions (Non-parametric Kruskal-Wallis test corrected with Dunn’s test for multiple comparisons; ns: non-significant; *p<0.05; **p<0.01; ***p<0.001; ****p<0.0001. See also **Supplementary Figure 4**). Scale bars: 1mm. D: Dorsal; L: left; R: right; Ctx: Cortex; Hpc: Hippocampus; Th: Thalamus; CPu: Caudate Putamen; HTh: Hypothalamus; S1BF: barrel-field primary somatosensory cortex; S2: Secondary somatosensory cortex; AuD: Dorsal auditory cortex; VPM: Ventral postero-medial nucleus of the thalamus; ; VPL: Ventral postero-lateral nucleus of the thalamus; Po: Posterior nucleus of the thalamus.

### Functional ultrasoung imaging acquisition

Coronal μDoppler images were acquired using a 15-MHz linear probe composed of 128 piezo-elements spaced by 100μm (L22-14Vx, Vermon, France) connected to a dedicated ultrasound scanner (Vantage 128, Verasonics, USA) and controlled by a high-performance computing workstation (fUSI-2, AUTC, Estonia). This configuration allowed to image the brain vasculature with a resolution of 100μm laterally, 110μm in depth, and 300μm in elevation^34^. The ultrasound sequence generated by the software is adapted from Macé et al.^31^ and Brunner, Grillet et al.^34^ Ultrafast images of the brain were generated using 5 tilted plane-waves (−6°, -3°, +0.5°, +3°, +6°). Each plane wave is repeated 6 times and the recorded echoes are averaged to increase the signal to noise ration. The 5 plane-wave images are added to create compound images at a frame rate of 500Hz. To obtain a single vascular image we acquired a set of 250 compound images in 0.5s, an extra 0.3s pause is included between each image to have some processing time to display the images for real-time monitoring of the experiment. The set of 250 compound images has a mixed information of blood and tissue signal. To extract the blood signal we apply a low pass filter (cutt off 15Hz) and an SVD filter that eliminates 20 singular values. This filter aims to select all the signal from blood moving with an axial velocity higher than ∼1mm/s. To obtain a vascular iimage we compute the intensity of the blood signal i.e., Power Doppler image. This image is in first approximation proportional to the cerebral blood volume^26,28^. Overall, this process enables a continious acquisition of power Doppler images at a frame rate of 1.25Hz during several hours. Then, the acquired images are processed with a dedicated GPU architecture, displayed in real-time for data visualization, and stored for subsequent off-line analysis^34^.

### fUS data processing and analysis

The data processing was performed following the procedure described by Brunner, Grillet et al.,^34^.

#### Registration to Paxinos rat brain atlas and data segmentation

We registered the fUS dataset to a custom digital rat brain atlas used in Brunner et al.^35^, using one coronal plane (bregma -3.4mm) from the stereotaxic atlas of Paxinos^42^. The image of the brain vasculature was manually translated and rotated to align it the coronal plane of the reference atlas. For an accurate registration, we used landmarks such as the surface of the brain, hippocampus, superior sagittal sinus, and other large vessels. If needed, the brain volume was scaled to fit the atlas outline. The outcome of this registration procedure is an affine coordinate transformation : 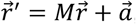, where 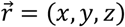 are the original coordinates image of the brain vasculature, M is the rotation and scaling matrix and 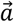 the translation vector. The dataset was segmented into 69 anatomical regions/hemispheres of the reference atlas (see **Supplementary Table 2**). The hemodynamic signals were averaged in each area. The segmentation and the data processing were performed using an automated MatLab-based pipeline. The software for data registration and segmentation is available in open-access^34^.

#### Relative cerebral blood volume (rCBV)

We used the relative cerebral blood volume (rCBV, expressed in % as compared to baseline) to analyze ischemia, transient hemodynamic events associated with spreading depolarizations (SDs) and functional changes. rCBV is defined as the signal in each voxel compared to its average level during the baseline period. After registration and segmentation, the rCBV signal was averaged in each individual region.

#### Analysis of stroke hemodynamics

The extraction of the temporal traces from the ischemic area was performed based on the temporal analysis of the rCBV signal in the primary somatosensory barrel-field cortex (S1BF). The detection of hemodynamic events associated with SDs was performed based on the temporal analysis of the rCBV signal in the retrosplenial granular (RSGc) and dysgranular (RSD) cortices of the left hemisphere (ipsilesional). Hemodynamic events associated with SDs were defined as transient increase of rCBV signal (+25%) detected with a temporal delay of <10 frames (i.e., 8s) between the two regions of interest, validating both the hyperemia and spreading features of hemodynamic events associated with spreading depolarizations^35,43–45^. This procedure allowed to measure the occurrence of hemodynamic event associated with SDs over the recording period. Live recording of ischemia and spreading depolarizations can be visualized in **Movie 1**.

#### Activity maps

Pre- and post-stroke recordings are reshaped in 40-s sessions, i.e., 50 frames, centered on the start of the stimulation (at 20s), and averaged based on the whisker stimulation paradigm (left or right). In each voxel, we compared signals along the recording in a time window before the stimulus onset and a time window after stimulus onset using two-tailed Wilcoxon rank sum test. We obtained the z-statistics of the test for each voxel, and consequently a z-score for the coronal cross-section. Mean activity maps for left or right whisker stimulation (**Figure 3B** and **4A**) show z-score value calculated using a Fisher’s transform for all voxels across the coronal cross-section. Only voxels with a z-score>1.6 were considered significantly activated (p<0.05 for a one-tailed test).

**Figure 4.**
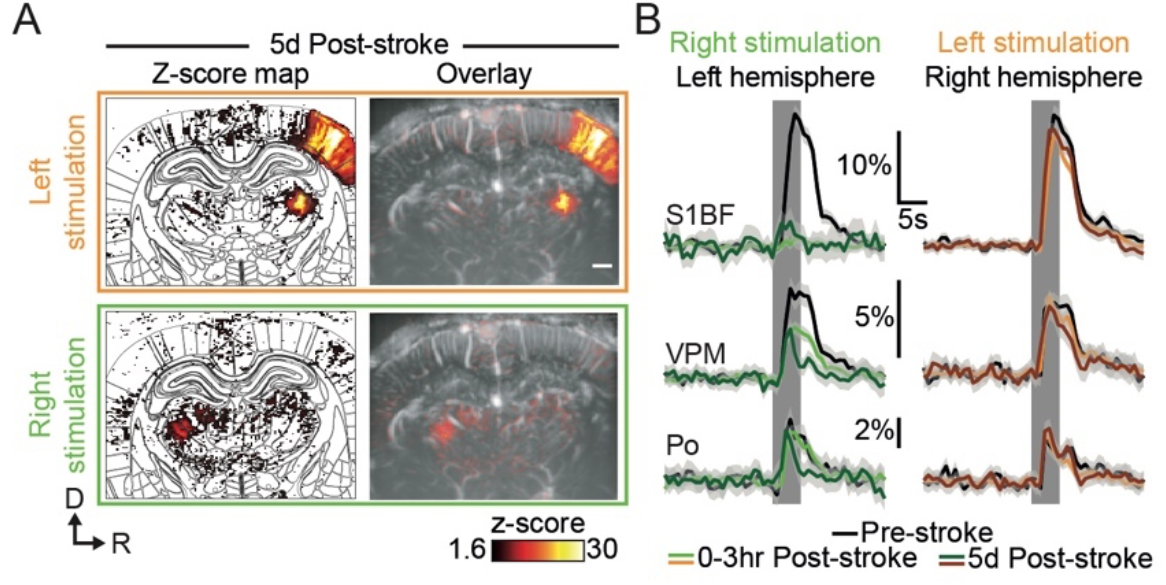
Late post-stroke alteration of whisker-to-barrel thalamo-cortical circuit. **(A)** Activity maps (z-score; average of 45 trials) depicting evoked functional responses to left (orange) or right whisker pads stimulation (green) 5d after stroke induction. Z-score maps are registered with the Paxinos atlas (white outlines; Left) and overlaid with the corresponding coronal μDoppler image (Right). **(B)** Left; Average response curves to left and right whisker stimulation (orange and green; respectively) extracted from S1BF, VPM, and Po before (Pre-stroke, black, n=2, 45 trials/rat), 0-3hr (0-3hr Post-Stroke; light orange/green, n=2, 45 trials/hr/rat) and 5d after stroke induction (5d Post-stroke, dark orange/green, n=2, 45 trials/rat). Data are mean±95%CI. The vertical bar represents the whisker stimulus. Scale bars: 1mm. D: Dorsal; R: right; S1BF: barrel-field primary somatosensory cortex; VPM: Ventral postero-medial nucleus of the thalamus; Po: Posterior nucleus of the thalamus.

#### Hemodynamic response time-courses

The relative hemodynamic time course ΔrCBV was computed for each brain regions (after registration and segmentation; **Figure 3C-D** and **4B**), as the rCBV change compared to baseline at each time point. No additional filtering was used, and no trial was removed from the analysis.

#### Statistical analysis

Activated brain regions were detected from hemodynamic response time-courses using GLM followed by t-test across animals as proposed in Brunner, Grillet et al.,^34^. The area under the curve (AUC) from hemodynamic response time-courses was computed for individual trials in S1BF, VPM and Po regions, for all the periods of the recording and for all rats included in this work. AUC were compared and analysed using a non-parametric Kruskal-Wallis test corrected for multiple comparison using a Dunn’s test. Tests were performed using GraphPad Prism 10.0.1.

### Histopathology

Rats were killed 24hrs after the occlusion for histological analysis of the infarcted tissue. Rats received a lethal injection of pentobarbital (100mg/kg i.p. Dolethal, Vetoquinol, France). Using a peristaltic pump, they were transcardially perfused with phosphate-buffered saline followed by 4% paraformaldehyde (Sigma-Aldrich, USA). Brains were collected and post-fixed overnight. 50-μm thick coronal brain sections across the MCA territory were sliced on a vibratome (VT1000S, Leica Microsystems, Germany) and analyzed using the cresyl violet (Electron Microscopy Sciences, USA) staining procedure (see <u>Open Lab Book</u> for procedure). Slices were mounted with DPX mounting medium (Sigma-Aldrich, USA) and scanned using a bright-field microscope.

## Results

### Animals

Report on animal use, experimentation, exclusion criteria can be found in **Supplementary Table 1**. Rat #1 was excluded after the control session as the imaging window was too anterior to capture both cortical and thalamic responses. Rat #2 was excluded as hemodynamic responses were inconsistent during baseline (pre-stroke) period. Rat #9 showed early post-stroke reperfusion and was excluded from stroke analysis, the control session (pre-stroke) from Rat #9 was analyzed.

### Real-time imaging of stroke induction in awake rats

We first developed a dedicated procedure for real-time imaging of stroke induction and associated evoked functional deficits in awake head-fixed rats (**Figure 1A**). Each rat was subjected to two cranial windows accessing independently the distal branch of the left MCA (**Figure 1B**, Left) and the selected brain regions to image (**Figure 1B**, Right). The latter was performed between bregma -2 and -4mm allowing for jointly monitoring the bilateral thalamo-cortical circuits of the somatosensory whisker-to-barrel pathway, including the ventroposterior medial nucleus of the thalamus (VPM) and the primary somatosensory barrel-field cortex (S1BF). Moreover, the selected coronal cross-section includes the posterior nucleus of the thalamus (Po), the reticular nucleus of the thalamus, and the ventral part of the zona incerta known for relaying information related to whiskers^46,47^, and also direct efferent projections from the S1BF to other cortical and subcortical regions^48^. Prior to imaging sessions, rats were extensively trained to accept comfortable restraints in the experimental apparatus (**Figure 1C**), suitable for fUS recording of brain functions and stroke induction under awake conditions. After data acquisition, the coronal cross-section was registered and segmented on a custom-developed digital rat atlas^49^ to provide a dynamic view of the changes in perfusion induced either by the stroke or evoked activity.

To overcome the limitations of conventional stroke models, we occluded the distal branch of the MCA by the mean of a chemo-thrombotic ferric chloride solution (FeCl_3_)^36,37^ while performing fUS imaging in awake rats (**Figure 2A**). It should be mentioned that the rats did not show any obvious signs of pain or discomfort (e.g., vocalization, aggressiveness) during the restrain period and occlusion procedure. The MCA occlusion (MCAo) was captured live with fUS and confirmed by the large drop of signal, i.e., ischemia, localized in the cortex of the left hemisphere (**Figure 2B, C, Movie 1** and **Supplementary Figure 1**) as shown with μDoppler image taken 3hrs and 5d after the stroke onset (dashed outline, **Figure 2B**, Top row). Bmode images accounting for the brain tissue echogenicity remain unchanged early after stroke onset (3hrs) while showing focal hyper-echogenicity (dashed outline, **Figure 2B**, Bottom row) lately after stroke onset (5d) as a marker of focal lesion^50^. The stroke-induced hemodynamic changes have been continuously recorded for up to 3hrs after stroke onset, registered and segmented into 69 regions (**Supplementary Figure 1**). We first extracted the average change in rCBV (ΔrCBV in %) in the S1BF cortex of the left hemisphere (blue region, **Figure 2B**) and detected an abrupt drop of rCBV down to ∼40% of the baseline level after the occlusion of the MCA, followed by a progressive decrease of the rCBV to 30% of baseline level 3hrs after the stroke onset (**Figure 2C** and **Movie 1**). Second, we extracted the average rCBV change from a cortical region supplied by the anterior cerebral artery directly after the MCAo. The signal extracted from the retrosplenial granular cortex (RSGc; purple and black regions in **Figure 2B**) shows successive and transient increases of signal. It characterizes hemodynamic events associated with spreading depolarizations (SDs) in the left hemisphere (in purple; **Figure 2D** and **Movie 1**) while resulting in a slight and stable oligemia in the right hemisphere (in black; **Figure 2D** and **Supplementary Figure 1**). SD events were observed in the peri-ischemic territory of all rats subjected to MCAo and occurred in an ostensibly random fashion (**Figure 2E**); however, hemodynamic events associated with SDs showed a similar bell shape and time-course across animals (**Figure 2F**). On average, we detected 5 SD events per hour per rat. Finally, we stained brain slices 24hrs after MCAo and confirmed that FeCl_3_-induced ischemia turned into tissue infarction (red delineation; **Figure 2G**).

### Stroke-induced alterations of the thalamo-cortical functions

One hour before and during 3hrs after the occlusion of the MCA, rats received mechanical stimulation of the whisker alternately delivered to the left and right pad using motorized combs (5-Hz sinusoidal deflection, 20° amplitude, 5-s duration; **Figure 3A**) to capture the spatiotemporal dynamics of the functional circuit. Before stroke, the sensory-evoked stimulations elicited a robust and statistically significant functional response (z-score>1.6, see Material and Methods) for both left and right stimulation (orange and green respectively; z-score maps; Pre-stroke panel, **Figure 3B** and **Movie 2**) with the activity spatially confined in the contralateral dorsal part of the VPM and S1BF. The temporal analysis of the somatosensory evoked responses in the contralateral hemisphere confirmed that VPM, Po, and S1BF regions were significantly activated and for both left and right stimuli (****p<0.0001, ***p<0.001 and ****p<0.0001 respectively; Left panel, **Figure 3C**). We also detected significant increase of activity in S2, AuD, Ect (****p<0.0001) and PRh (***p<0.001) cortices and VPL nucleus (**p<0.01; the list of acronyms is provided in **Supplementary Table 2**), brain regions receiving direct efferent projections from the S1BF^48,51,52^, VPM or Po nuclei^53–55^. It is worth noted that no habituation or sensitization due to the repetitiveness of whiskers stimulation was observed in cortical and subcortical regions over the pre-stroke sessions (**Supplementary Figure 2**).

After the stroke, the activity map from the left pad stimulation elicited a similar response pattern as pre-stroke; however, the right pad stimulation showed a total absence of functional response in the S1BF cortex and a significant reduction of the response in the VPM (z-score maps; Post-stroke panel, **Figure 3B** and **Movie 2**). Over the 3hrs folowing stroke onset, functional responses to left whisker stimulation were still detected in the cortical and thalamic regions of the contralateral (right) hemisphere; however, functional responses to right whisker stimulation were only detected in subcortical nuclei (i.e., VPM, Po, VPL), while attenuated when compared with the responses from the pre-stroke period and from the other hemisphere (**Figure 3B, C**). Furthermore, no responses were detected at the cortical level (S1BF, S2, and AuD; right panel, **Figure 3B, C**). A larger version of **Figure 3C** is provided in **Supplementary Figure 3**.

To better evaluate how the functional responses were affected by the stroke, we have divided the post-stroke recording period into 3 sections of 1hr each and compared them with the 1-hr pre-stroke period (**Figure 3D**). Temporal plots from the pre-stroke period showed robust increases in signal during the stimulus in S1BF, VPM, and Po regions and high consistency between left and right stimuli (black plots, **Figure 3D** and **Supplementary Figures 2-3**); fitting well the hemodynamic response functions as previously observe^16,56^. Indeed, the hemodynamic responses were characterized by a quick increase in signal during whisker stimulation reaching a peak after 4.0s at 18.2±1.3% (4.0s, 18.6±1.2%) of baseline level for S1BF, 4.0s at 4.6±0.5% (3.2s, 5.8±0.7%) for VPM, and 2.4s at 2.9±0.7% (3.2s, 4.0±0.8%) for Po from the left stimulation (right, respectively; mean±95%CI) before slowly returning to baseline level (black plots, **Figure 3D**).

During the first hour following the stroke onset, functional responses in the left hemisphere (i.e., ipsilesional) were abolished in the S1BF, S2, and AuD (0-1hr Post-stroke, ****pvalue<0.0001), significantly decreased in the VPM (0-1hr Post-stroke, ***pvalue<0.001), and unchanged in Po and VPL (0-1hr Post-stroke, ns; **Figure 3D**) when compared with the pre-stroke responses (Pre-stroke, black plots, **Figure 3D**). Over the two following hours (i.e., 1-2hr and 2-3hr Post-stroke), the hemodynamic responses captured in these regions remained similar as those detected during the first post-stroke hour (green plots, **Figure 3D**).

Regarding the right hemisphere (i.e., contralesional), the functional responses of S1BF and VPM were conserved during the first hour after the stroke onset (ns, 0-1hr Post-stroke; orange plots, **Figure 3D**). During the two following hours, signal changes in S1BF show a significant and progressive decrease of activity (1-2hr Post-stroke **pvalue<0.01, 2-3hr Post-stroke ****pvalue<0.0001; orange plots, **Figure 3D**; Similar observation were made for S2 and AuD) whereas responses in VPM remained stable during the second hour post-stroke (1-2hr, ns) before significantly decreasing during the third hour (2-3hr Post-stroke *pvalue<0.05; orange plots, **Figure 3D**). Finally, the functional responses in VPM and Po remained unchanged over the 3hrs following the stroke onset (bottom panel, **Figure 3D**).

Activity maps, region-time traces of the 69 brain regions, mean and individual time-course for all trials (left and right stimuli - including ipsi and contralateral traces), imaging timepoints (Control, Pre-Stroke, Post-Stroke) for all the rats included in this work can be found in **Supplementary Figure 5**.

### Delayed alteration of the somatosensory thalamo-cortical pathway

A secondary objective of this work was to evaluate the fUS ability to identify potential delayed functional alteration within a few days after the initial injury. Two animals were imaged five days after the MCAo following the same experimental, stimulation, imaging, and processing conditions as for the early post-stroke session. As the number of rats imaged at this timepoint is limited to n=2, we did not performed statistical analysis. Activity maps, region-time traces and individual trials for both right and lieft stimulation (including ipsi and contralateral) for the each rats are provided in **Supplementary Figure 5**.

Five days after the MCA occlusion, we first placed the ultrasound probe over the imaging window and adjusted its position (using micromanipulator) to find back the recording plane from Pre-Stroke session using Bmode (morphological mode) and μDoppler imaging using brain vascular landmarks (i.e., vascular patterns, brain surface and hippocampus^34,35^; see **Figure 2B**). Functional responses to left whisker stimulation were still detected in the right hemisphere (i.e., contralesional), at the cortical and subcortical levels (orange; **Figure 4A**). As for the early post-stroke imaging period, the functional responses to right whisker stimulation were only detected in the subcortical nuclei and not at the cortical level (green; **Figure 4A**).

Second, we extracted and compared the temporal plots of the functional responses gathered 5d after the stroke with the one obtained from the same two animals at the pre-stroke and 3hrs post-stroke timepoints (**Figure 4B**). At this later time point, the functional responses in the left S1BF (dark green plot, left panel, **Figure 4B**. Similar observation were made for the S2 and AuD) remained abolished when compared with the pre-stroke period (black plot), while slightly increased when compared with the 3hrs post-stroke timepoint (green plot). The responses detected in the VPM 5d after the stroke onset (dark green plot, left panel, **Figure 4B**) were largely decreased not only when compared with the pre-stroke signal (black plot) but also with the 3hrs post-stroke trace (green plot). Interestingly, both the amplitude and time-to-peak of the hemodynamic response function were very similar to those from the early post-stroke signal (i.e., 3hrs post-stroke); however, the post-peak period was largely dampened in the 5d post-stroke signal. A similar alteration of the hemodynamic response function was also observed for the 5d post-stroke signal extracted from the Po nucleus when compared to the pre-stroke and 3hrs post-stroke signals (left panel, **Figure 4B**. Similar observation were made for the VPL).

Regarding the right hemisphere (i.e., non-ischemic; right panel, **Figure 4B**), the S1BF functional responses to left whisker stimulation were still reduced when compared with pre-stroke responses (black plot) but remained similar to the traces detected at 3-hr post-stroke (orange plot, non-significant). As for the left VPM, both the amplitude and time-to-peak of the hemodynamic responses from the right VPM responses were consistent with pre-stroke and 3hr post-stroke values but the post-peak signal was decreased (brown plot). The functional responses extracted from the Po and VPL did not show changes when compared to pre-stroke and 3hrs post-stroke responses.

## Discussion

With this proof-of-concept study, we document on the feasibility of the continuous brain hemodynamics recording of a focal cerebral ischemia after MCAo in conscious rats. Using functional ultrasound imaging, we were able to extract multiple parameters (i.e., ischemia, location and spreading depolarization), characteristic of such cortical stroke. Then, we report on how the functional sensorimotor thalamo-cortical circuit was altered at early and late post-stroke stages.

Compared to highly-invasive conventional strategies such as clipping or suturing^1,2^, the FeCl_3_ model used here, is well suited to study stroke under awake conditions. Indeed, the use of FeCl_3_ requires less manipulation, allows to maintain the dura intact and strongly reduces the risk of hemorrhage^36,37^ and animal loss. Furthermore, the FeCl_3_ model closely mimics key human stroke features including focal ischemia, creation of blood clot, possibility of vessel recanalization, and penumbral tissue^36,37^. However, to adequatly and efficiently occlude the vessel of interest, removing a piece of skull remains required. As mentioned in the report on animal use, one rat was excluded from the analysis as the MCA spontaneously reperfuses, thus dropping the success rate of such model.

The FeCl_3_-induced MCAo showed an abrupt and massive drop of blood perfusion remaining constant during the entire recording period. The ischemia was confined within the cortical territory perfused by the MCA (**Figure 2B**), and the infarct (location and size; **Figure 2G**) is in agreement with previous observations^37,49^. We also detected transient hyperemic events associated with spreading depolarizations (SDs) within the peri-ischemic territory, with occurence, frequency and amplitude of the hemodynamic waves (**Figure 2D-F**) consistent with prior observation^43,49,57–61^. Moreover, the spatiotemporal dynamic of the FeCl_3_-induced MCAo is consistent with previous fUS imaging reports on cortical ischemia with various stroke model^16,49,62^.

On top of tracking large hemodynamic variation (i.e. ischemia, SDs), one asset of the fUS imaging technology relies on its ability to track subtle hemodynamic changes in sparse brain regions^16,26,29,31,33,34,63^. Therefore, we have evaluated how evoked functional responses reorganize at early and late timepoints after stroke induction. Functional responses to mechanical whisker stimulation were detected in several regions relaying the information from the whisker to the cortex, including the VPM and Po nuclei of the thalamus, and S1BF, the somatosensory barrel-field cortex. Responses were also observed in the S2 cortex involved in the multisensory integration of the information^46,47,64^, the auditory cortex as it receives direct efferent projection from S1BF^48,64^, and the VPL nuclei of the thalamus connected via corticothalamic projections^48^

Functional responses extracted in the left hemisphere affected by the focal ischemia (i.e., ipsilesional) show a primary alteration of the whisker-to-barrel pathway within the first hour after the stroke onset. While the abrupt loss in S1BF responses was mainly driven by the focal ischemia, the immediate but partial drop in VPM responses (**Figure 3D**) might result from the direct the loss of the excitatory corticothalamic feedback to the VPM^55,65,66^, or even from a dampening of thalamocortical excitability^67^. The absence of such cortical feedback suggests that the dampened functional responses might be driven by the intrinsic activity of the VPM in response to whisker stimulation. Five days after the initial injury, nuclei of the thalamus (VPM and Po) were subjected to a delayed and robust functional alteration (**Figure 4B**) as previously confirmed in other thalamic relay^68^, probably associated with diaschisis, as previously characterised by tissue staining, reduction of metabolism, functions and perfusion^21–23,53,68^. Functional responses of the S1BF extracted from the right hemisphere (i.e., contralesionel) show a significant decrease shortly after the stroke onset (**Figure 3D**), and still detected at day 5, could be provoked by a loss of transcortical excitability^69,70^. The late drop in VPM responses might be explained by corticothalamic modulation of the projections toward VP^46,70^.

While preliminary, these results obtained from awake head-fixed rats are in contradiction with a similar work by our group (fUS imaging, distal MCAo with microvascular clip, electrical whisker stimulation) showing higher contralesional responses to whisker stimulation during early stages of ischemic stroke^16^. However, these experiments were subjected to long-term isoflurane regimen (surgery and imaging) known to alter functional responses^71–73^ as well as disrupting hemodynamics^40^. Therefore, further studies will be needed to accurately dissect the complex and long-lasting post-stroke alterations of the functional whisker-to-barrel pathway, including at the neuronal level by direct electrophysiology recordings and imaging, as fUS only reports on hemodynamics as a proxy of local neuronal activity^30,31,63,74–76^. Another limitation relies on the experimental condition as our brain imaging paradigm was constrained to a single cross-section, thus missing out-of-plane brain regions also affected by the stroke (e.g., ischemic size, infract extension, origin, and propagation pattern of SDs)^41^ or involved in the whisker network (e.g., superior colliculus, striatum, amygdala and cerebellum)^46^. To overcome such limitation, one can extend the size of the cranial window to allow for larger scale imaging either by sequentially scanning the brain^30,31,34,35,62,75,77,78^, or by using the recently developed volumetric fUS which provides whole-brain imaging capabilities in anesthetized^79^ and awake rats^33^. Finally, it is important to note that this proof-of-concept work did not specifically focus the impact of sex dimorphism on the stroke or early behavioral outcomes following the insult that would greatly enhance the translational value of such preclinical stroke study^80^.

Beyond studying the whisker-to-barrel somatosensory circuit, the brain-wide capability of fUS opens the door to investigate on stroke-affected brain circuits and functions using transgenic lines combined with opto-/chemo-genetic strategies as the technology is fully mature for mice studies^31,33,34,75^.

## Supplementary Materials

**Supplementary Table 1.** Reporting on animal use, experimentation, exclusion criteria, and figure association.

| Animal Tag | Imaging session |  |  |  |  | Issue |
| --- | --- | --- | --- | --- | --- | --- |
|  | Control | Baseline (1hr) | Stroke | Early Post-Stroke (3hrs) | Late Post-Stroke (1hr) |  |
| Rat #1 | Session excluded | Not performed |  |  | Not performed | Miss functional responses from thalamic nuclei as the imaging plane was too anterior |
| Rat #2 | Not performed | Session excluded |  |  | Not performed | Unconsistency of functional responses during baseline. Issue with cranial window |
| Rat #3 | Not performed | Session Included |  |  | Not performed |  |
| Rat #4 | Not performed | Session Included |  |  | Not performed |  |
| Rat #5 | Session Included | Not performed |  |  | Not performed |  |
| Rat #6 | Not performed | Session Included |  |  | + 5d<br>Session Included |  |
| Rat #7 | Not performed | Session Included |  |  | Not performed |  |
| Rat #8 | 2d before stroke<br>Session Included | Session Included |  |  | + 5d<br>Session Included |  |
| Rat #9 | 2d before stroke<br>Session Included | Session excluded |  |  | Not performed | Reperfusion of the MCA after stroke onset |
| Sessions used in | - Supplementary<br>Figures 2, 4, 5 | - Figures 2 and 3<br>- Movies 1 and 2<br>- Supplementary Figures 1 to 5 |  |  | - Figure 4<br>- Supplementary<br>Figures 4 and 5 |  |

**Supplementary Table 2.**
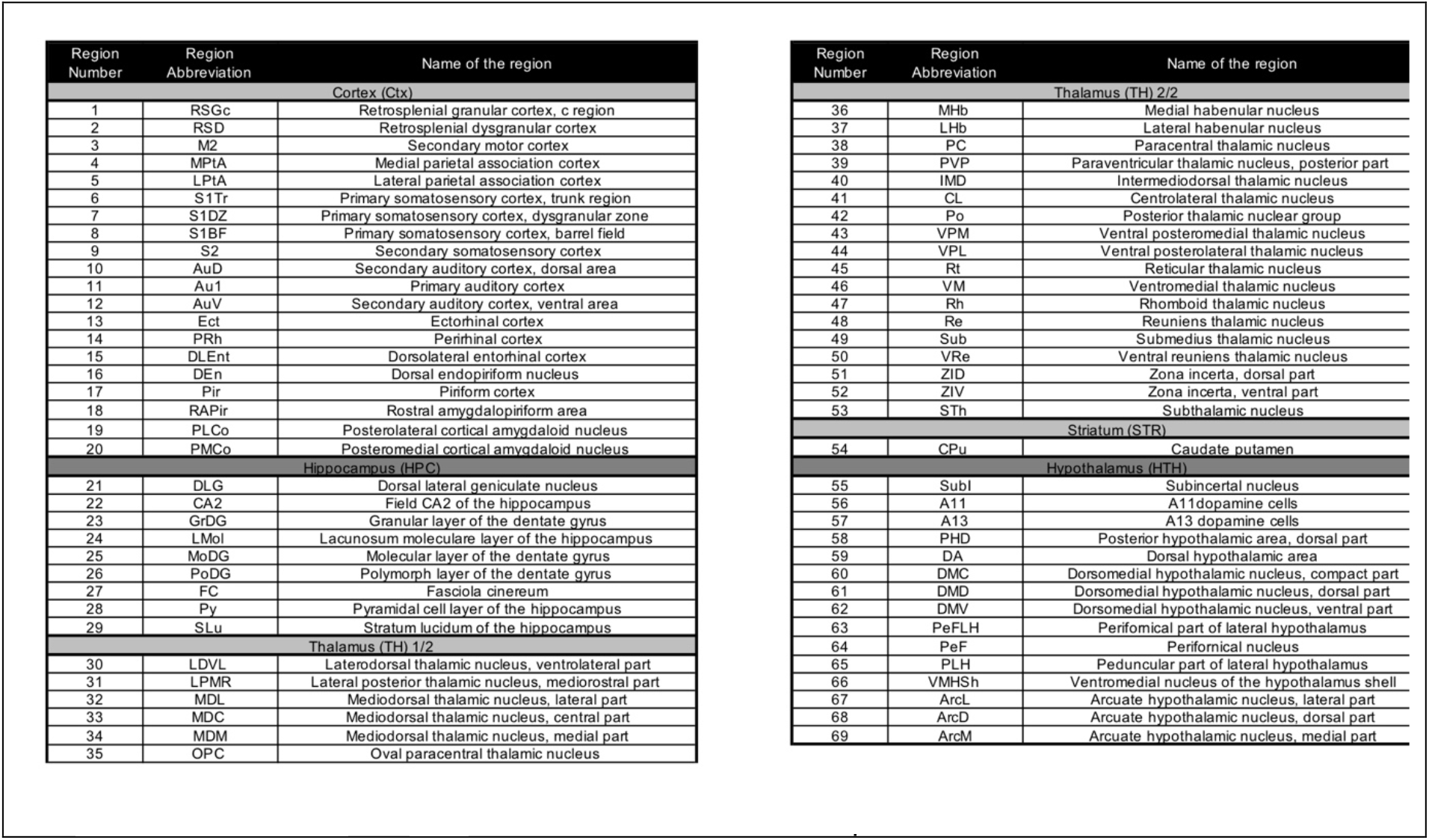
List of the 69 brain regions/hemisphere from the coronal cross-section μDoppler imaged in each rat organized by main anatomical structures. Adapted from the Paxinos rat brain atlas^42^.

**Movie 1**. Movie of hemodynamic changes induced by MCA occlusion using FeCl_3_ in awake head-fixed rats. Raw images.

**Movie 2**. Movie of thalamo-cortical functional responses to left and right whisker stimulation before and 3hrs after stroke onset.

**Supplementary Figure 1.**
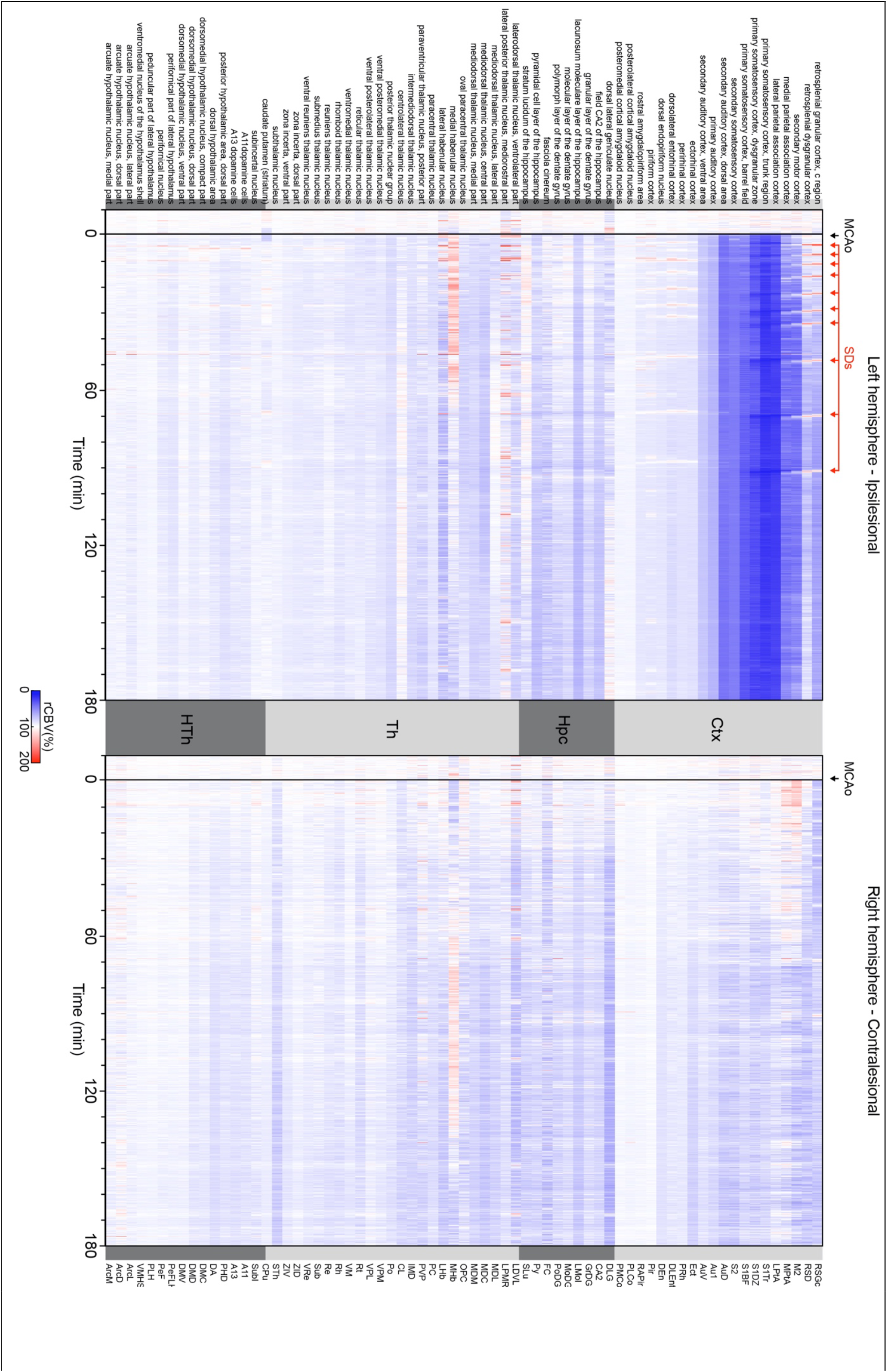
Hemodynamic changes (rCBV in %) induced by MCAo in 69 regions located in the ipsilesional (left panel) and contralesional hemisphere (right panel) of the imaged coronal cross-section. Regions are organized by main anatomical structures (see **Supplementary Table 2**). SDs stands for hemodynamic events associated with spreading depolarizations.

**Supplementary Figure 2.**
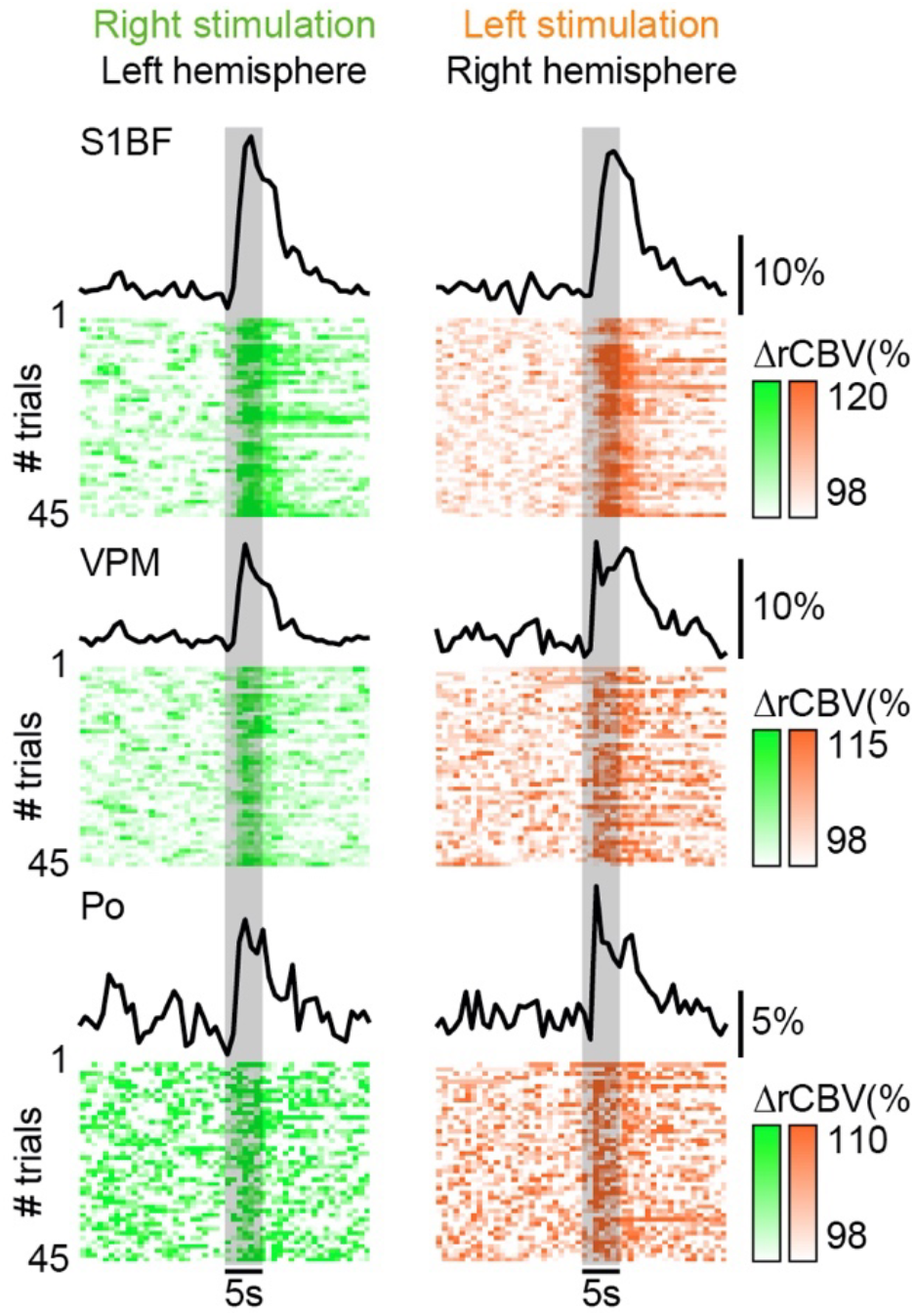
Averaged hemodynamic response curves (ΔrCBV in %) of 45 consecutive right (green) or left whisker stimulation (orange; 1hr recording) extracted in the contralateral S1BF, VPM and Po regions (top to bottom). The corresponding individual trials presented below confirmed the stability across the recording. Vertical grey bar, the period of whisker stimulation.

**Supplementary Figure 3.**
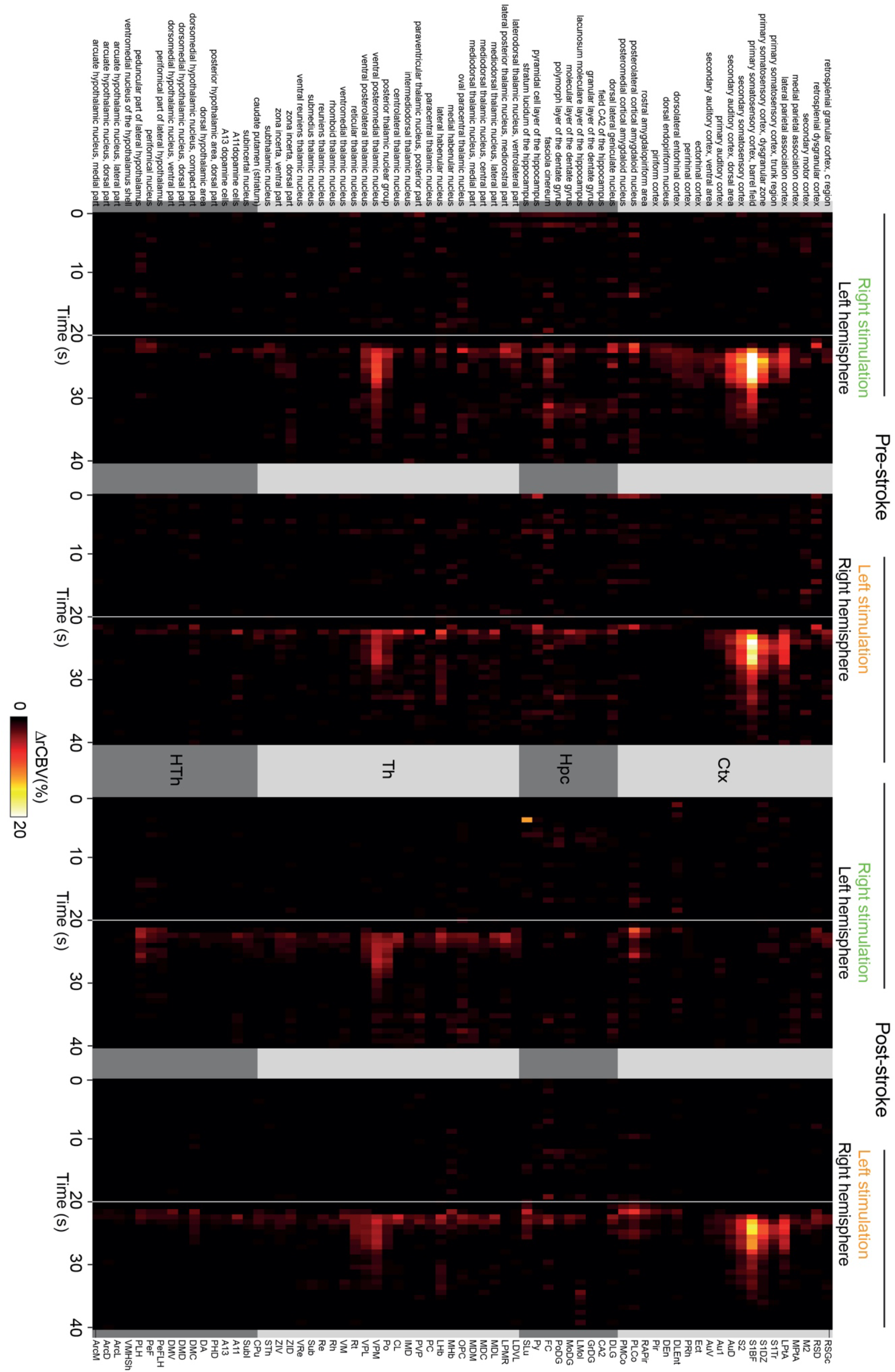
Close up view of Figure 3C.

**Supplementary Figure 4.**
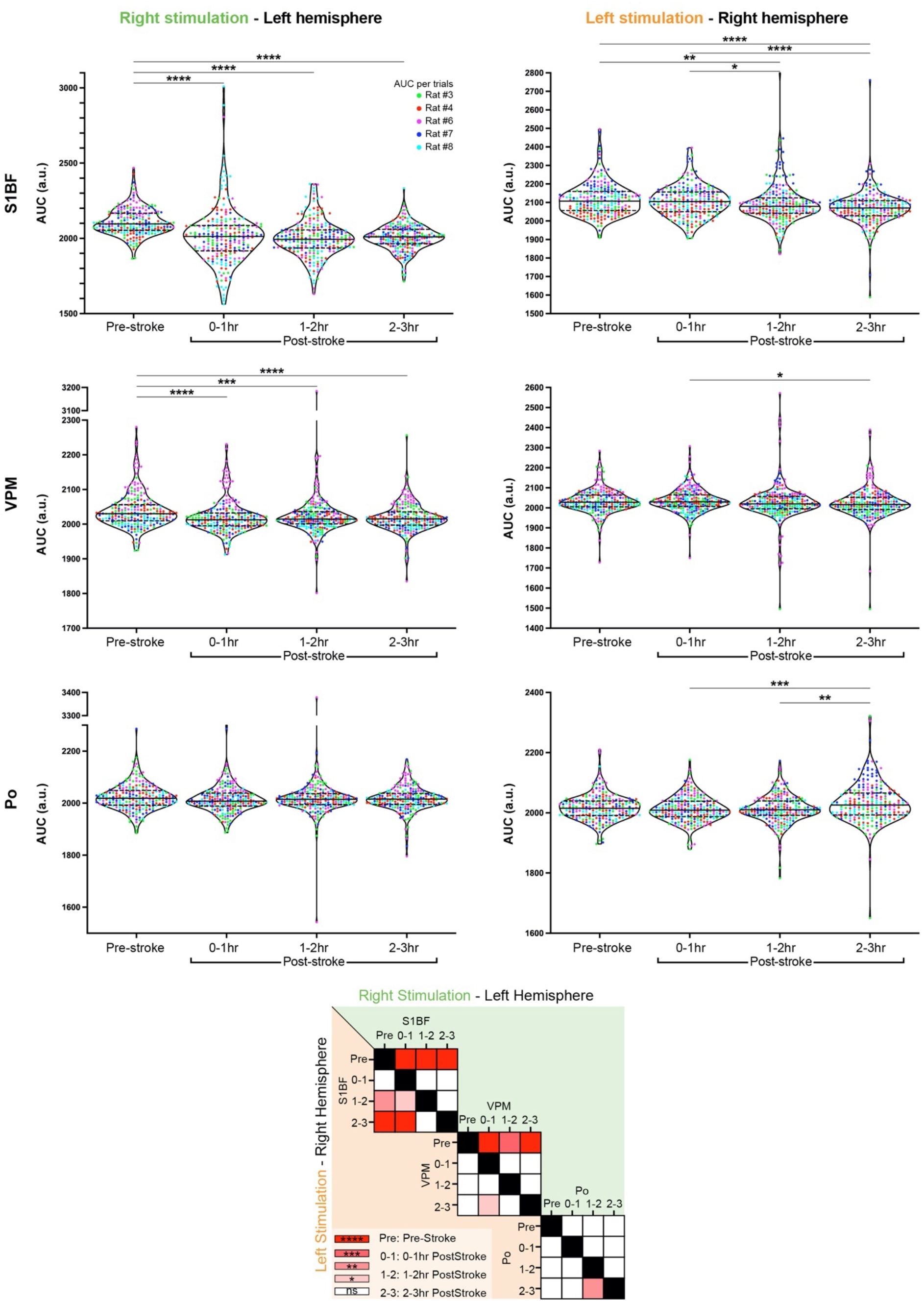
**Top Panel -** Violin plots showing the distribution of the area under the curve (AUC) extracted from hemodynamic response time-courses of individual trials in S1BF (top row), VPM (middle row) and Po regions (bottom row), for stimulation delivered either to the right (left column) or left whisker pad (right column) along all the periods of the recording (Pre-Stroke, 0-1hr Post-stroke, 1-2hr Post-Stroke, 2-3hr Post-Stroke). Each dot represents an inidivudal trial, each color represent a rat. **Bottom Panel –** Matrix comparing AUC from S1BF, VPM, and Po for right (green - top right diagonal) or left stimulation (orange - bottom left diagonal) at Pre-Stroke, 0-1hr Post-stroke, 1-2hr Post-Stroke, and 2-3hr Post-Stroke timepoints. AUC were compared and analysed using a non-parametric Kruskal-Wallis test corrected for multiple comparison using a Dunn’s test.

**Supplementary Figure 5.**
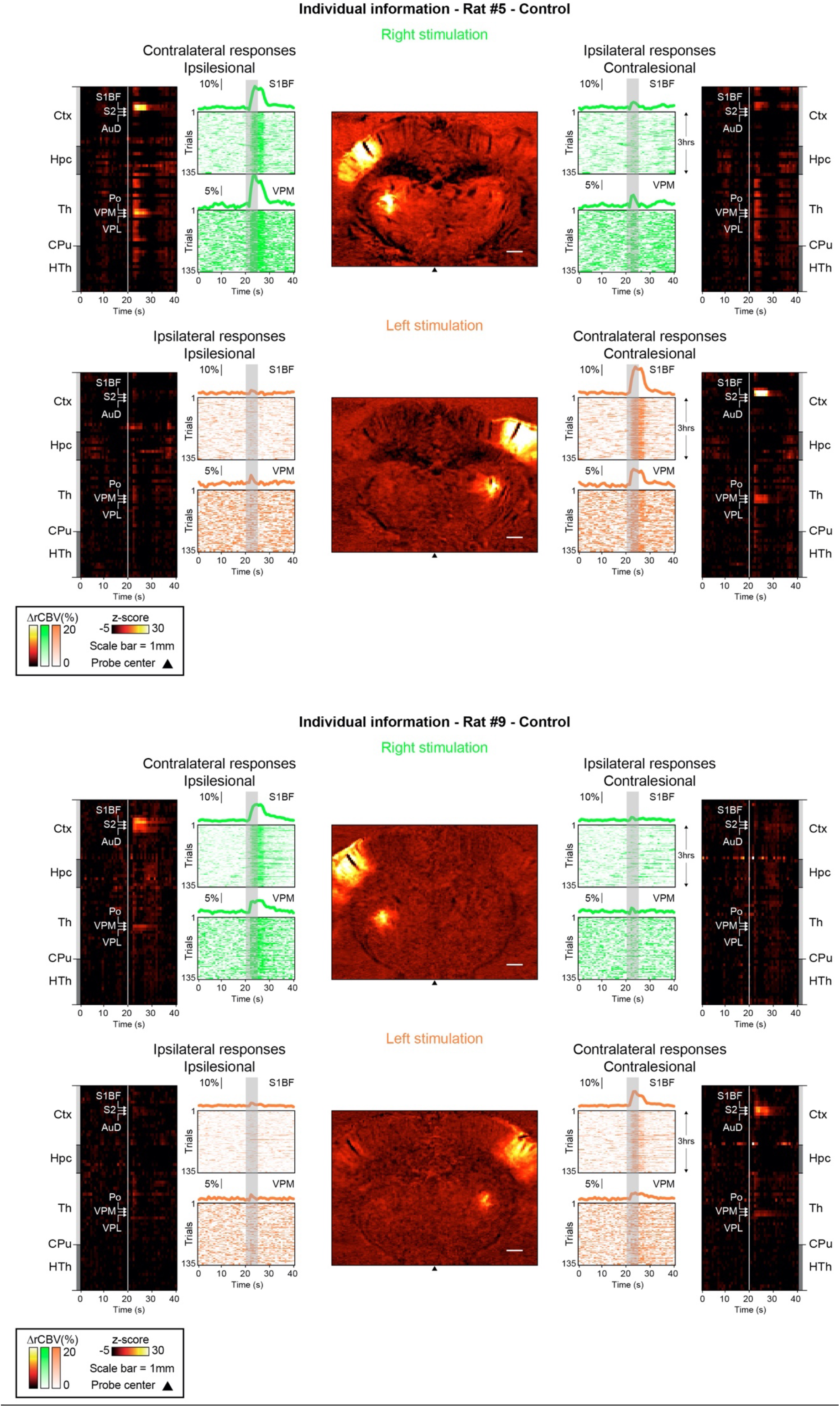
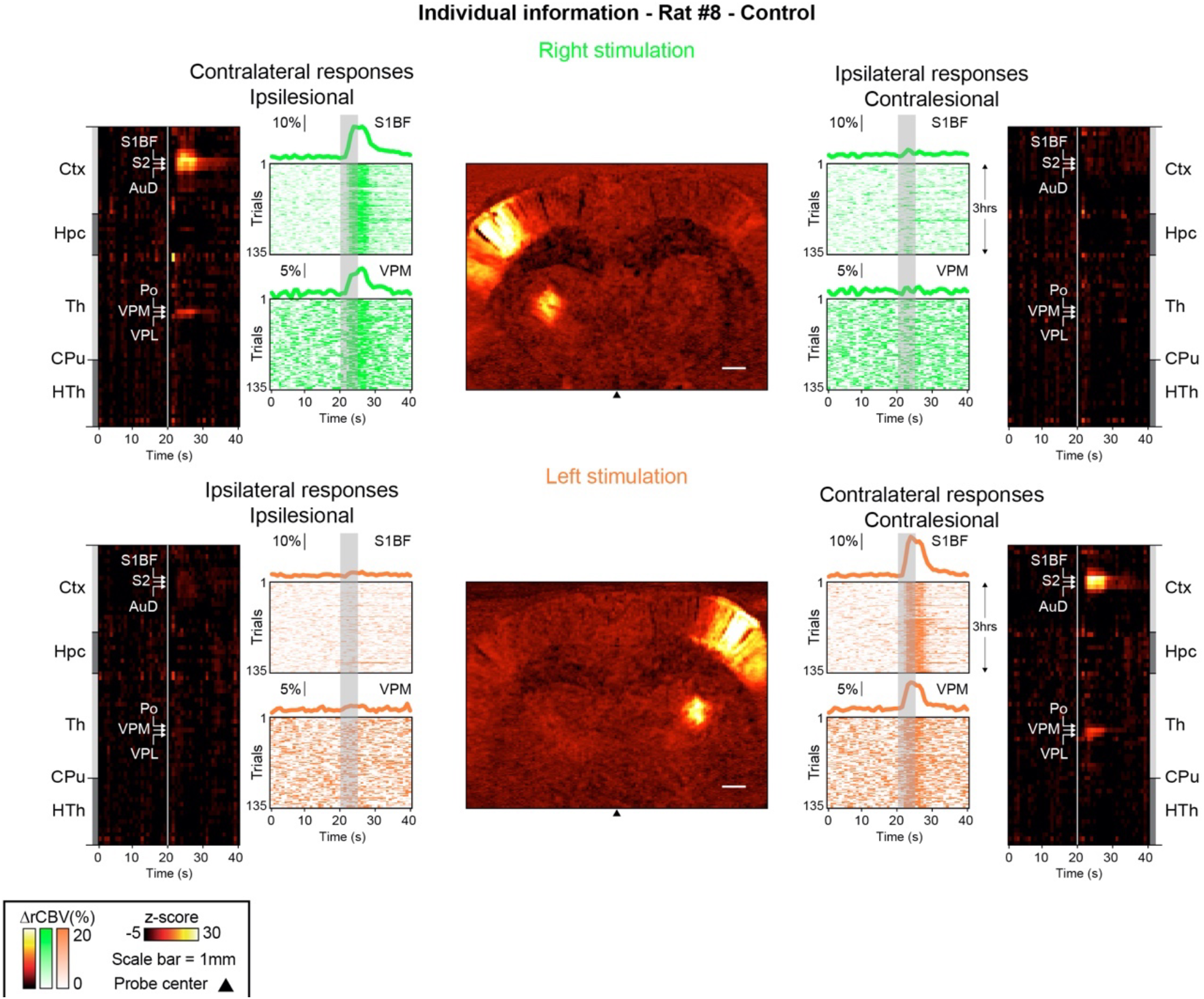
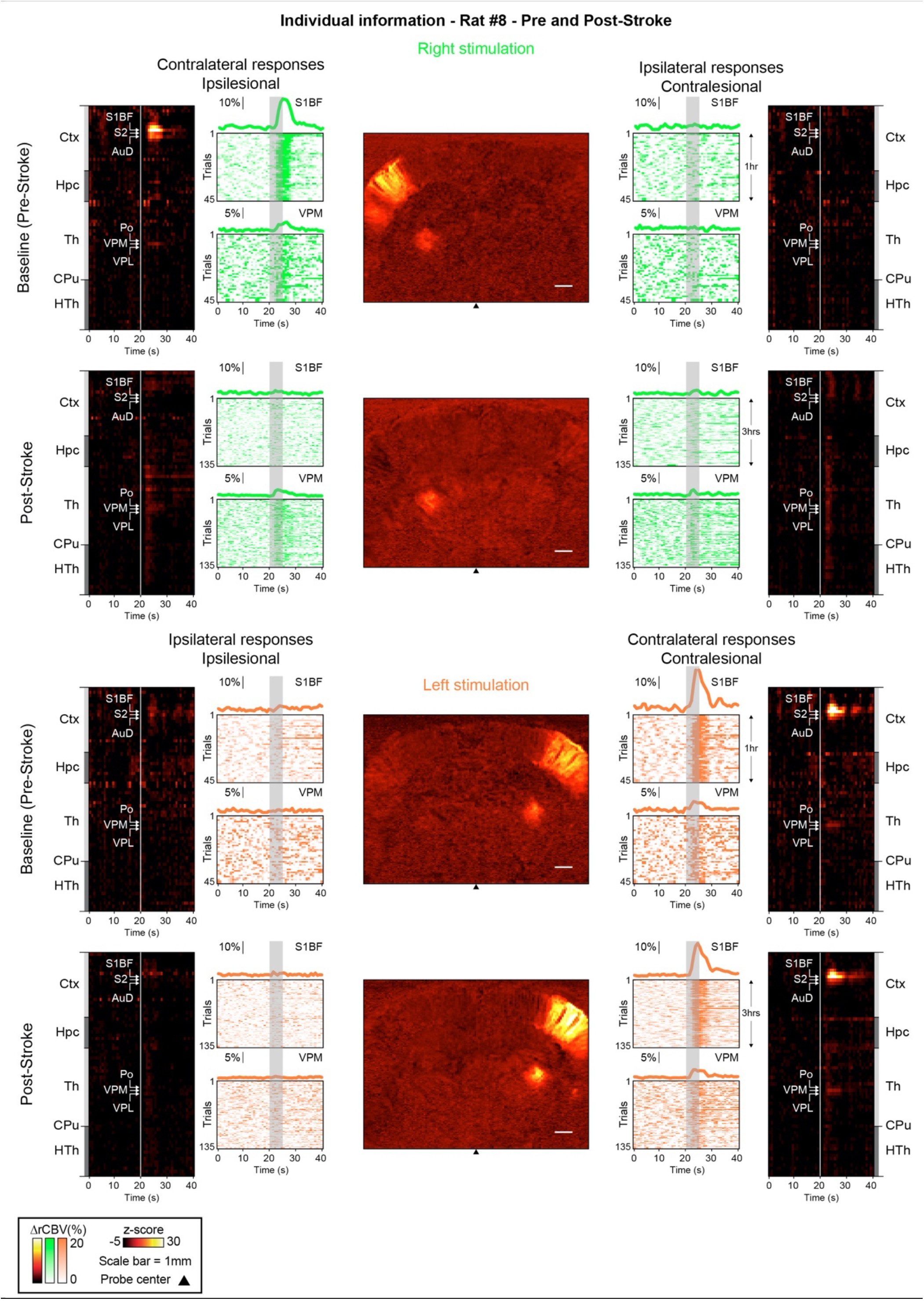
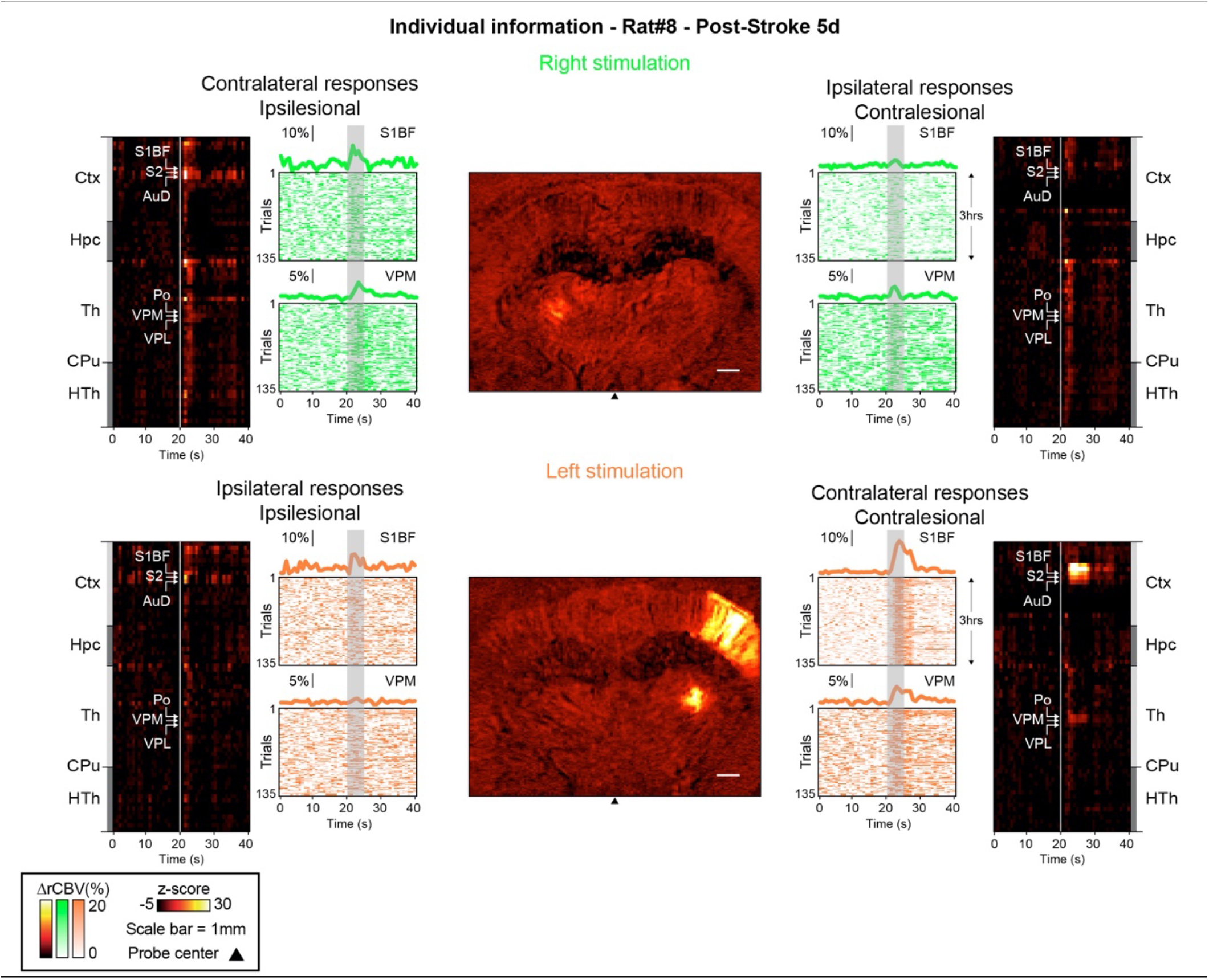
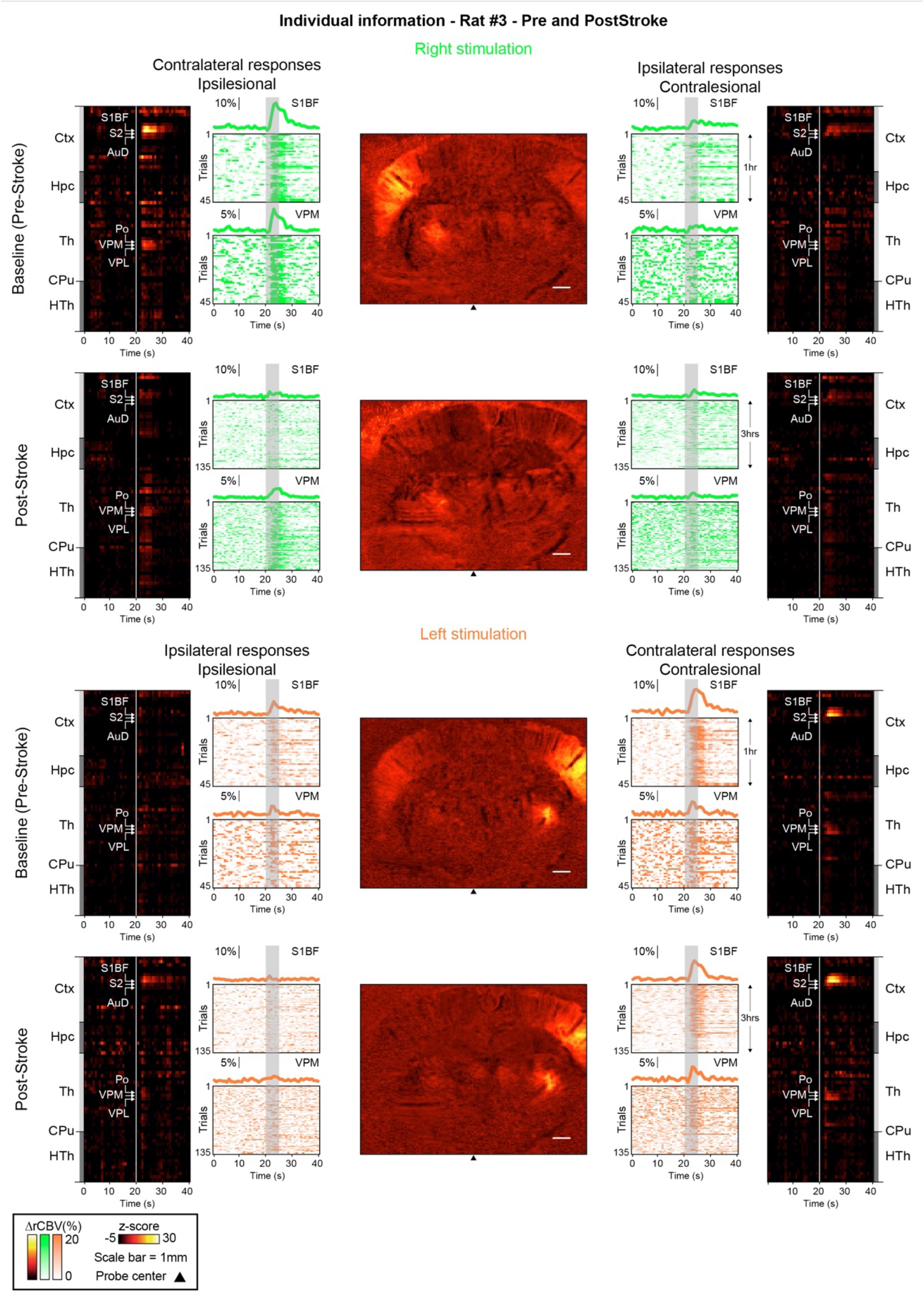
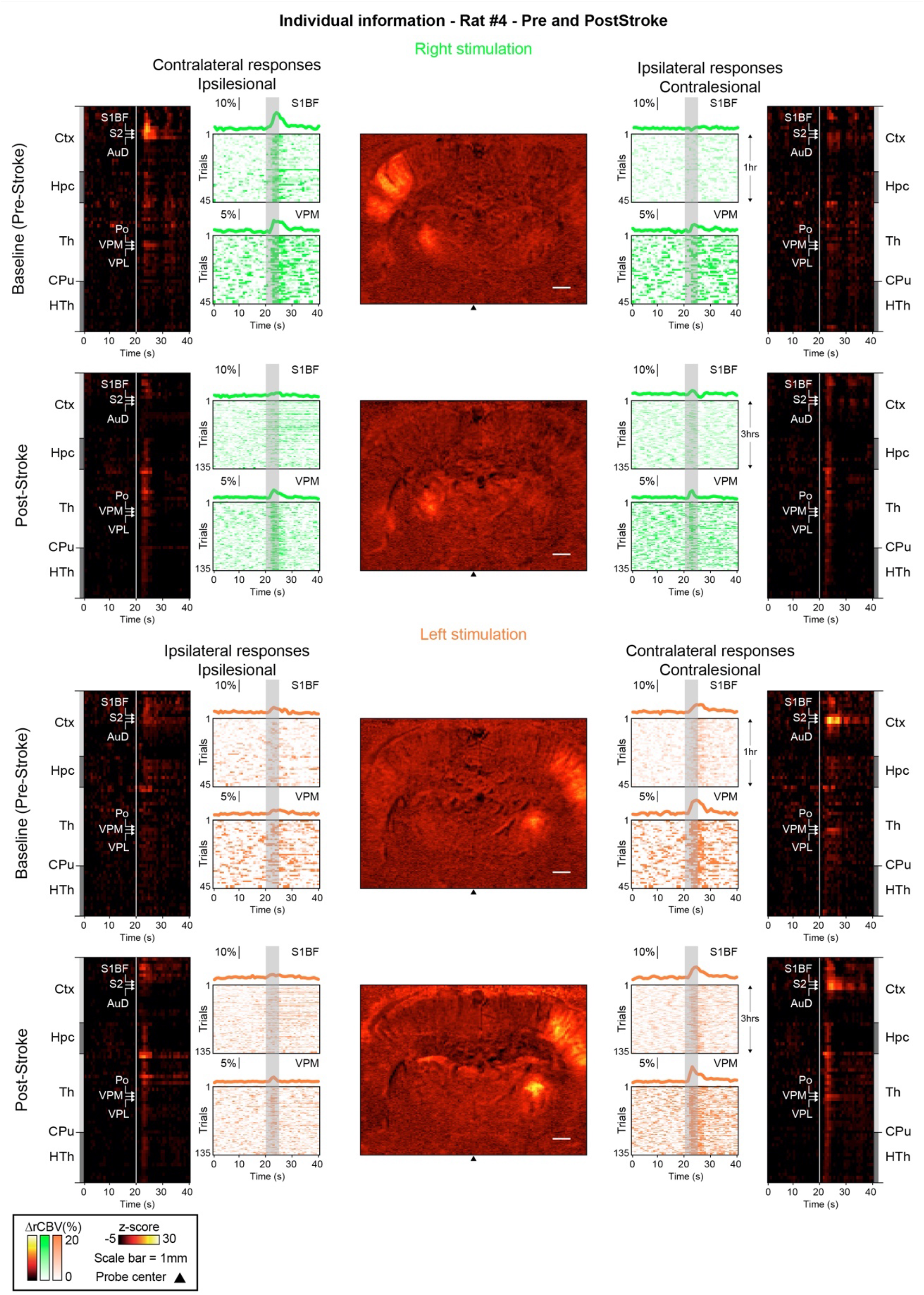
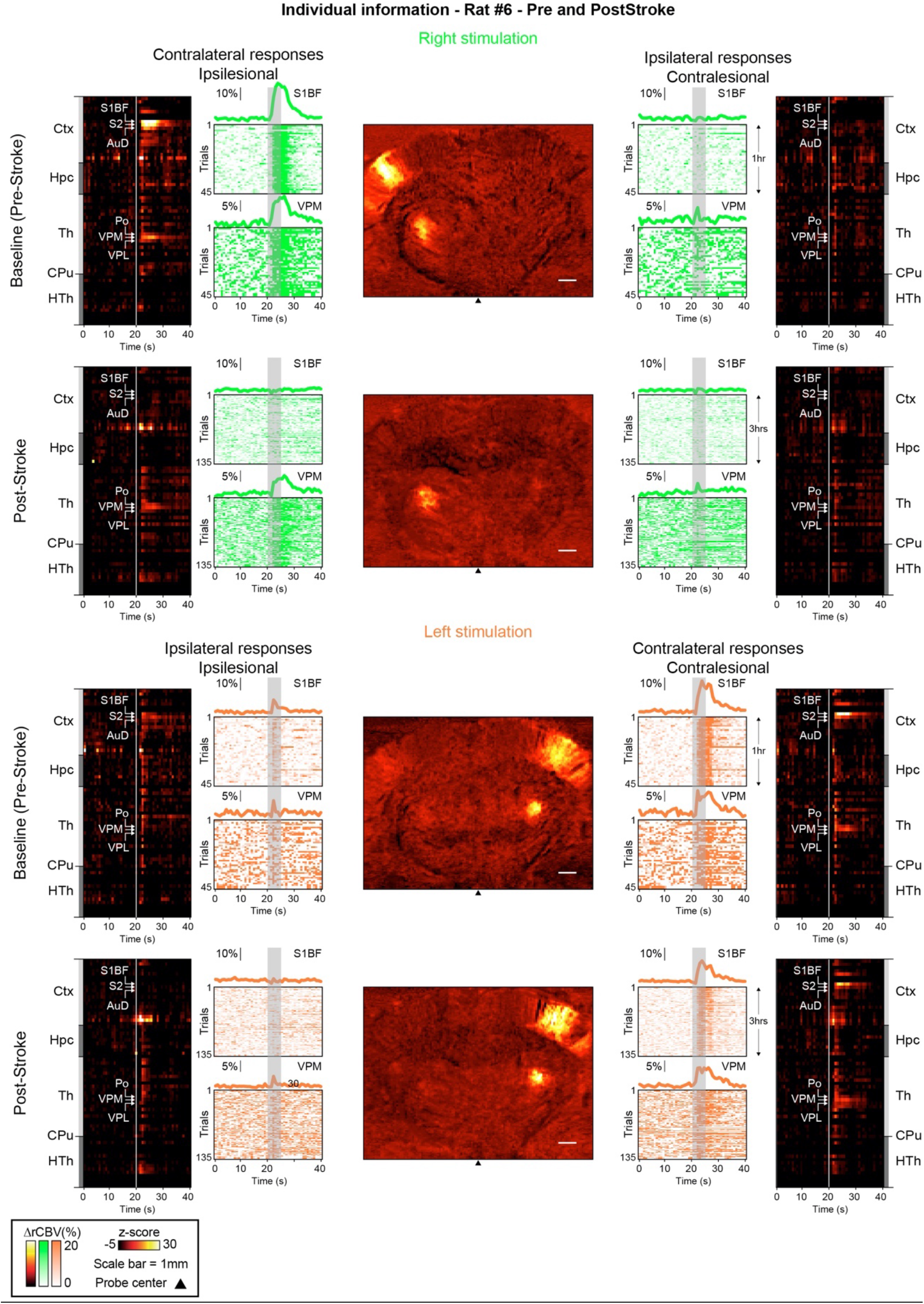
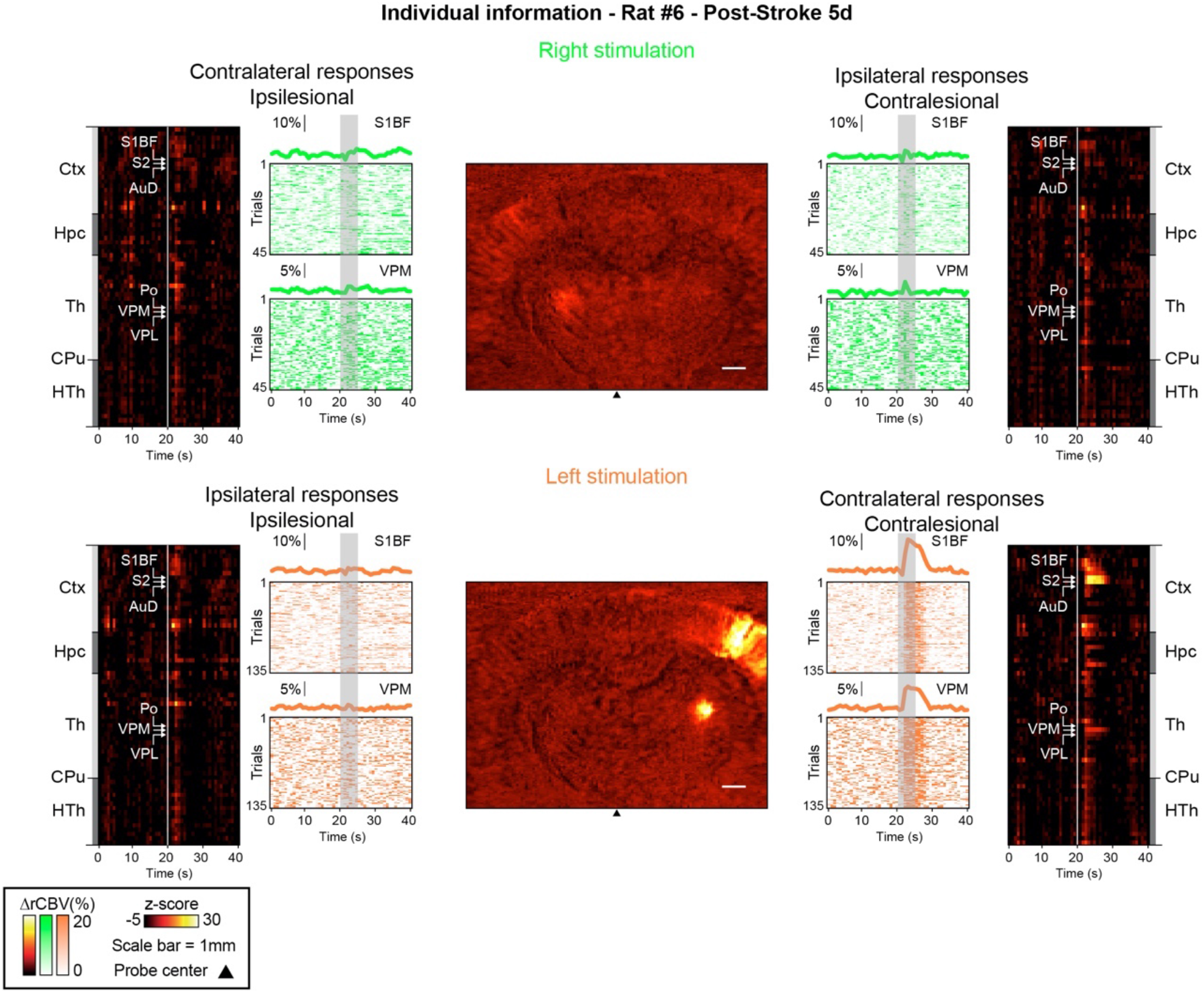
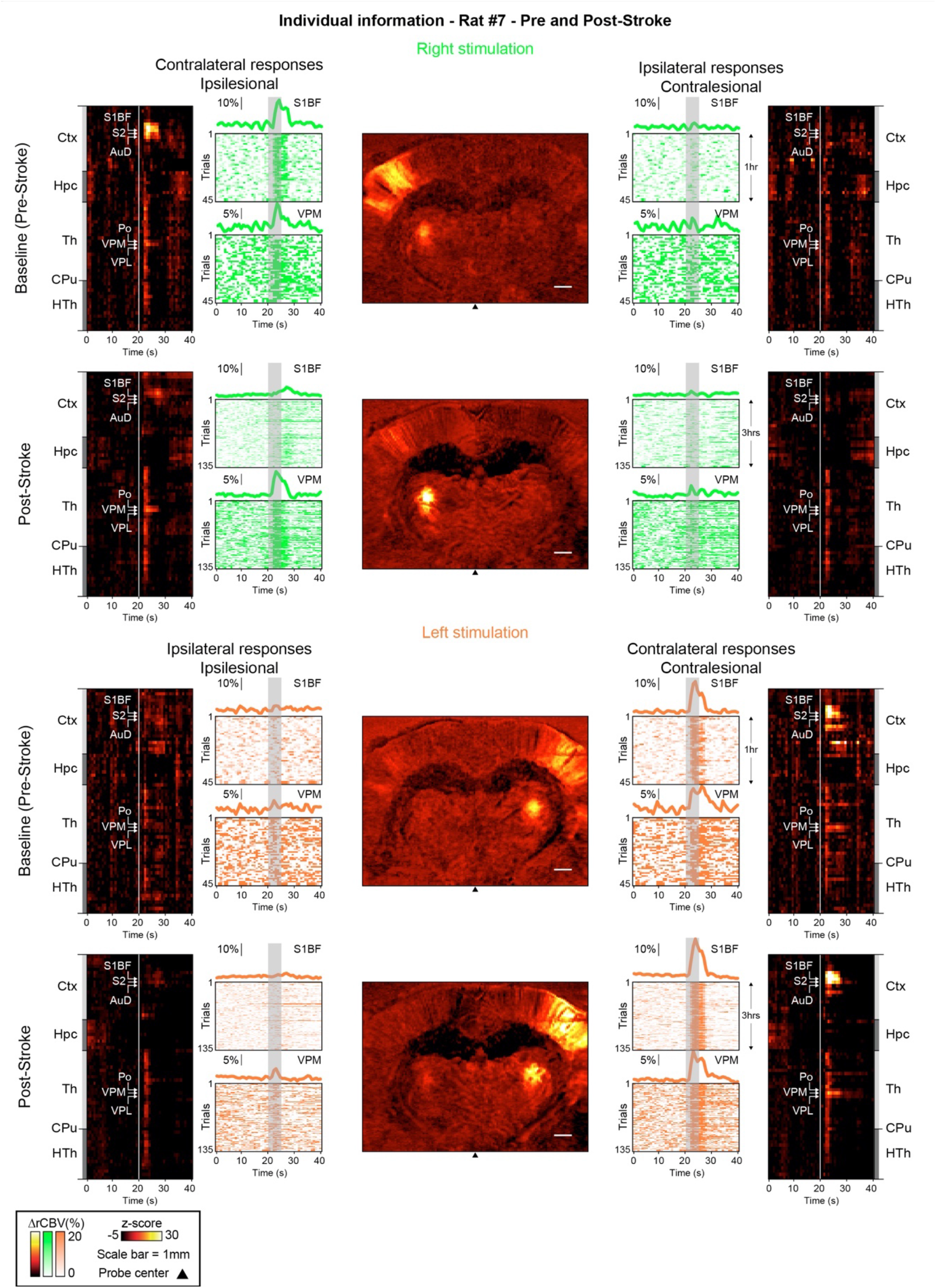
Activity maps, region-time traces of the 69 brain regions imaged, mean and individual time-courses for all trials (left and right stimuli - including contra- and ipsilateral traces) and imaging timepoints (Control, Pre-Stroke, Post-Stroke) for all the rats included in this work.

## Author contribution

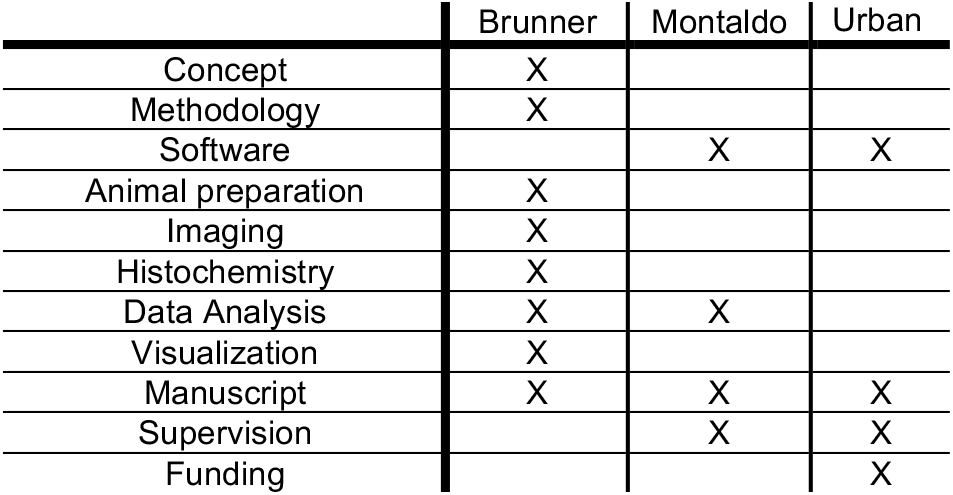

## Funding

This work is supported by grants from the Fondation Leducq (15CVD02) and KU Leuven (C14/18/099-STYMULATE-STROKE). The functional ultrasound imaging platform is supported by grants from FWO (MEDI-RESCU2-AKUL/17/049, G091719N, and 1197818N), VIB Tech-Watch (fUSI-MICE), Neuro-Electronics Research Flanders TechDev fund (3D-fUSI project).

## Acknowledgements

The authors thank the members of the Fondation Leducq network #15CVD02, Dr. M. Grillet, T. Lambert and lab members for their insightful comments and discussions. We thank NERF animal caretakers, including I. Eyckmans, F. Ooms, and S. Luijten, for their help with the management of the animals. Figures 1-3 use BioRender.com icons.

## Competing interests

A.U. is the founder and a shareholder of AUTC company commercializing functional ultrasound imaging solutions for preclinical and clinical research.

## Notes

### Competing Interest Statement

Alan URBAN is the founder and a shareholder of A.U.T.C. LLC, a technology consulting company.

### Summary of Updates

We have thoroughly reviewed the feedback provided by the reviewers and addressed each of their inquiries and apprehensions. In line with their suggestions, we have revised our manuscript to incorporate greater details regarding the experimental procedures and the statistical methodologies employed. Moreover, we have eliminated certain analyses that were based on a limited number of imaging sessions. Given that our paper is categorized as a Resource and Tools article, we have appropriately recalibrated the assertion of our study to align with its nature as a proof-of-concept paper, which presents preliminary findings that are robust and dependable. Concurrently, we have introduced three new Supplementary Figures, among which one exhibits data obtained from individual animals collectively.

## References

1. MacRae, I. Preclinical stroke research -Advantages and disadvantages of the most common rodent models of focal ischaemia. Br. J. Pharmacol. 164, 1062–1078 (2011).

2. Fluri, F., Schuhmann, M. K. & Kleinschnitz, C. Animal models of ischemic stroke and their application in clinical research. Drug Des. Devel. Ther. 9, 3445–3454 (2015).

3. Sommer, C. J. Ischemic stroke: experimental models and reality. Acta Neuropathologica vol. 133 245–261 Preprint at https://doi.org/10.1007/s00401-017-1667-0 (2017).

4. Mackey, J. et al. Population-based study of wake-up strokes. Neurology 76, 1662–1667 (2011).

5. Muir, K. W. Treatment of wake-up stroke: stick or TWIST? Lancet Neurol. 22, 102–103 (2023).

6. Reimann, H. M. & Niendorf, T. The (Un)Conscious Mouse as a Model for Human Brain Functions: Key Principles of Anesthesia and Their Impact on Translational Neuroimaging. Front. Syst. Neurosci. 14, 1–42 (2020).

7. Masamoto, K. & Kanno, I. Anesthesia and the Quantitative Evaluation of Neurovascular Coupling. J. Cereb. Blood Flow Metab. 32, 1233–1247 (2012).

8. Traystman, R. J. Effect of anesthesia in stroke models. in Neuromethods 121–138 (Humana Press, 2010).

9. Hoffmann, U., Sheng, H., Ayata, C. & Warner, D. S. Anesthesia in Experimental Stroke Research. Transl. Stroke Res. 7, 358–367 (2016).

10. Slupe, A. M. & Kirsch, J. R. Effects of anesthesia on cerebral blood flow, metabolism, and neuroprotection. J. Cereb. Blood Flow Metab. 38, 2192–2208 (2018).

11. Seto, A. et al. Induction of ischemic stroke in awake freely moving mice reveals that isoflurane anesthesia can mask the benefits of a neuroprotection therapy. Front. Neuroenergetics 6, p(2014).

12. Lu, H. et al. Induction and imaging of photothrombotic stroke in conscious and freely moving rats. J. Biomed. Opt. 19, 096013 (2014).

13. Balbi, M. et al. Targeted ischemic stroke induction and mesoscopic imaging assessment of blood flow and ischemic depolarization in awake mice. Neurophotonics 4, 035001 (2017).

14. Sunil, S. et al. Awake chronic mouse model of targeted pial vessel occlusion via photothrombosis. Neurophotonics 7, 015005 (2020).

15. Mohajerani, M. H., Aminoltejari, K. & Murphy, T. H. Targeted mini-strokes produce changes in interhemispheric sensory signal processing that are indicative of disinhibition within minutes. Proc. Natl. Acad. Sci. U. S. A. 108, E183–91 (2011).

16. Brunner, C. et al. Evidence from functional ultrasound imaging of enhanced contralesional microvascular response to somatosensory stimulation in acute middle cerebral artery occlusion/reperfusion in rats: A marker of ultra-early network reorganization? J. Cereb. Blood Flow Metab. 38, 1690–1700 (2018).

17. Dijkhuizen, R. M. et al. Functional magnetic resonance imaging of reorganization in rat brain after stroke. Proc. Natl. Acad. Sci. U. S. A. 98, 12766–12771 (2001).

18. Dijkhuizen, R. M. et al. Correlation between brain reorganization, ischemic damage, and neurologic status after transient focal cerebral ischemia in rats: a functional magnetic resonance imaging study. J. Neurosci. 23, 510–517 (2003).

19. Abo, M., Chen, Z., Lai, L. J., Reese, T. & Bjelke, B. Functional recovery after brain lesion--contralateral neuromodulation: an fMRI study. Neuroreport 12, 1543–1547 (2001).

20. Weber, R. et al. Early prediction of functional recovery after experimental stroke: functional magnetic resonance imaging, electrophysiology, and behavioral testing in rats. J. Neurosci. 28, 1022–1029 (2008).

21. Zhang, J., Zhang, Y., Xing, S., Liang, Z. & Zeng, J. Secondary neurodegeneration in remote regions after focal cerebral infarction: a new target for stroke management? Stroke 43, 1700–1705 (2012).

22. Carrera, E. & Tononi, G. Diaschisis: past, present, future. Brain 137, 2408–2422 (2014).

23. Cao, Z., Harvey, S. S., Bliss, T. M., Cheng, M. Y. & Steinberg, G. K. Inflammatory responses in the secondary thalamic injury after cortical ischemic stroke. Front. Neurol. 11, 236 (2020).

24. Levy, H., Ringuette, D. & Levi, O. Rapid monitoring of cerebral ischemia dynamics using laser-based optical imaging of blood oxygenation and flow. Biomed. Opt. Express 3, 777 (2012).

25. Dunn, A. K. Laser Speckle Contrast Imaging of Cerebral Blood Flow. Ann. Biomed. Eng. 40, 367–377 (2012).

26. Macé, E. et al. Functional ultrasound imaging of the brain. Nature methods 8, 662–664 (2011).

27. Demené, C., Mairesse, J., Baranger, J., Tanter, M. & Baud, O. Ultrafast Doppler for neonatal brain imaging. Neuroimage 185, 851–856 (2019).

28. Montaldo, G., Urban, A. & Macé, E. Functional ultrasound neuroimaging. Annu. Rev. Neurosci. 45, 491–513 (2022).

29. Urban, A. et al. Real-time imaging of brain activity in freely moving rats using functional ultrasound. Nature methods 12, 873–878 (2015).

30. Sieu, L.-A. et al. EEG and functional ultrasound imaging in mobile rats. Nat. Methods 12, 831–834 (2015).

31. Macé, É. et al. Whole-Brain Functional Ultrasound Imaging Reveals Brain Modules for Visuomotor Integration. Neuron 100, 1241–1251.e7 (2018).

32. Bergel, A. et al. Adaptive modulation of brain hemodynamics across stereotyped running episodes. Nat. Commun. 11, 6193 (2020).

33. Brunner, C. et al. A Platform for Brain-wide Volumetric Functional Ultrasound Imaging and Analysis of Circuit Dynamics in Awake Mice. Neuron (2020) doi:10.1016/j.neuron.2020.09.020.

34. Brunner, C. et al. Whole-brain functional ultrasound imaging in awake head-fixed mice. Nat. Protoc. (2021) doi:10.1038/s41596-021-00548-8.

35. Brunner, C. et al. Brain-wide continuous functional ultrasound imaging for real-time monitoring of hemodynamics during ischemic stroke. J. Cereb. Blood Flow Metab. 271678X231191600 (2023).

36. Karatas, H. et al. Thrombotic distal middle cerebral artery occlusion produced by topical FeCl(3) application: a novel model suitable for intravital microscopy and thrombolysis studies. J. Cereb. Blood Flow Metab. 31, 1452–1460 (2011).

37. Syeara, N. et al. Motor deficit in the mouse ferric chloride-induced distal middle cerebral artery occlusion model of stroke. Behav. Brain Res. 380, 112418 (2020).

38. Percie du Sert, N. et al. The ARRIVE guidelines 2.0: Updated guidelines for reporting animal research. PLoS Biol. 18, e3000410 (2020).

39. Martin, C. et al. Optical imaging spectroscopy in the unanaesthetised rat. J. Neurosci. Methods 120, 25–34 (2002).

40. Martin, C., Martindale, J., Berwick, J. & Mayhew, J. Investigating neural–hemodynamic coupling and the hemodynamic response function in the awake rat. Neuroimage 32, 33–48 (2006).

41. Topchiy, I. A. et al. Conditioned lick behavior and evoked responses using whisker twitches in head restrained rats. Behav. Brain Res. 197, 16–23 (2009).

42. Paxinos, G. The rat brain in stereotaxic coordinates. (Academic Press, 2014).

43. Bere, Z., Obrenovitch, T. P., Kozák, G., Bari, F. & Farkas, E. Imaging reveals the focal area of spreading depolarizations and a variety of hemodynamic responses in a rat microembolic stroke model. J. Cereb. Blood Flow Metab. 34, 1695–1705 (2014).

44. Ayata, C. & Lauritzen, M. Spreading depression, spreading depolarizations, and the cerebral vasculature. Physiol. Rev. 95, 953–993 (2015).

45. Binder, N. F. et al. Vascular response to spreading depolarization predicts stroke outcome. Stroke 53, 1386–1395 (2022).

46. Adibi, M. Whisker-mediated touch system in rodents: From neuron to behavior. Front. Syst. Neurosci. 13, 40 (2019).

47. Bosman, L. W. J. et al. Anatomical pathways involved in generating and sensing rhythmic whisker movements. Front. Integr. Neurosci. 5, 53 (2011).

48. Zakiewicz, I. M., Bjaalie, J. G. & Leergaard, T. B. Brain-wide map of efferent projections from rat barrel cortex. Front. Neuroinform. 8, 5 (2014).

49. Brunner, C. et al. Brain-wide continuous functional ultrasound imaging for real-time monitoring of hemodynamics during ischemic stroke. (2022) doi:10.1101/2022.01.19.476904.

50. Gómez-de Frutos, M. C. et al. B-mode ultrasound, a reliable tool for monitoring experimental intracerebral hemorrhage. Front. Neurol. 12, 771402 (2021).

51. Fabri, M. & Burton, H. Ipsilateral cortical connections of primary somatic sensory cortex in rats. J. Comp. Neurol. 311, 405–424 (1991).

52. Frostig, R. D., Xiong, Y., Chen-Bee, C. H., Kvasnák, E. & Stehberg, J. Large-scale organization of rat sensorimotor cortex based on a motif of large activation spreads. J. Neurosci. 28, 13274–13284 (2008).

53. Viaene, A. N., Petrof, I. & Sherman, S. M. Properties of the thalamic projection from the posterior medial nucleus to primary and secondary somatosensory cortices in the mouse. Proc. Natl. Acad. Sci. U. S. A. 108, 18156–18161 (2011).

54. El-Boustani, S. et al. Anatomically and functionally distinct thalamocortical inputs to primary and secondary mouse whisker somatosensory cortices. Nat. Commun. 11, 3342 (2020).

55. Landisman, C. E. & Connors, B. W. VPM and PoM nuclei of the rat somatosensory thalamus: intrinsic neuronal properties and corticothalamic feedback. Cereb. Cortex 17, 2853–2865 (2007).

56. Hirano, Y., Stefanovic, B. & Silva, A. C. Spatiotemporal evolution of the functional magnetic resonance imaging response to ultrashort stimuli. J. Neurosci. 31, 1440–1447 (2011).

57. Nakamura, H. et al. Spreading depolarizations cycle around and enlarge focal ischaemic brain lesions. Brain 133, 1994–2006 (2010).

58. Takeda, Y., Zhao, L., Jacewicz, M., Pulsinelli, W. A. & Nowak, T. S., Jr. Metabolic and perfusion responses to recurrent peri-infarct depolarization during focal ischemia in the Spontaneously Hypertensive Rat: dominant contribution of sporadic CBF decrements to infarct expansion. J. Cereb. Blood Flow Metab. 31, 1863–1873 (2011).

59. Farkas, E., Pratt, R., Sengpiel, F. & Obrenovitch, T. P. Direct, live imaging of cortical spreading depression and anoxic depolarisation using a fluorescent, voltage-sensitive dye. J. Cereb. Blood Flow Metab. 28, 251–262 (2008).

60. Farkas, E., Bari, F. & Obrenovitch, T. P. Multi-modal imaging of anoxic depolarization and hemodynamic changes induced by cardiac arrest in the rat cerebral cortex. Neuroimage 51, 734–742 (2010).

61. Bogdanov, V. B. et al. Susceptibility of primary sensory cortex to spreading depolarizations. J. Neurosci. 36, 4733–4743 (2016).

62. Hingot, V. et al. Early Ultrafast Ultrasound Imaging of Cerebral Perfusion correlates with Ischemic Stroke outcomes and responses to treatment in Mice. Theranostics 10, 7480–7491 (2020).

63. Urban, A. et al. Chronic assessment of cerebral hemodynamics during rat forepaw electrical stimulation using functional ultrasound imaging. Neuroimage 101, 138–149 (2014).

64. Lohse, M., Dahmen, J. C., Bajo, V. M. & King, A. J. Subcortical circuits mediate communication between primary sensory cortical areas in mice. Nat. Commun. 12, 3916 (2021).

65. Bourassa, J., Pinault, D. & Deschênes, M. Corticothalamic projections from the cortical barrel field to the somatosensory thalamus in rats: A single-fibre study using biocytin as an anterograde tracer. Eur. J. Neurosci. 7, 19–30 (1995).

66. Temereanca, S. & Simons, D. J. Functional topography of corticothalamic feedback enhances thalamic spatial response tuning in the somatosensory whisker/barrel system. Neuron 41, 639–651 (2004).

67. Tennant, K. A., Taylor, S. L., White, E. R. & Brown, C. E. Optogenetic rewiring of thalamocortical circuits to restore function in the stroke injured brain. Nat. Commun. 8, 15879 (2017).

68. Tokuno, T. et al. Functional changes in thalamic relay neurons after focal cerebral infarct: A study of unit recordings from VPL neurons after MCA occlusion in rats. J. Cereb. Blood Flow Metab. 12, 954–961 (1992).

69. Rema, V. & Ebner, F. F. Lesions of mature barrel field cortex interfere with sensory processing and plasticity in connected areas of the contralateral hemisphere. J. Neurosci. 23, 10378–10387 (2003).

70. Li, L., Rema, V. & Ebner, F. F. Chronic suppression of activity in barrel field cortex downregulates sensory responses in contralateral barrel field cortex. J. Neurophysiol. 94, 3342–3356 (2005).

71. Sicard, K. M., Henninger, N., Fisher, M., Duong, T. Q. & Ferris, C. F. Long-term changes of functional MRI-based brain function, behavioral status, and histopathology after transient focal cerebral ischemia in rats. Stroke 37, 2593–2600 (2006).

72. Paasonen, J. et al. Comparison of seven different anesthesia protocols for nicotine pharmacologic magnetic resonance imaging in rat. Eur. Neuropsychopharmacol. 26, 518–531 (2016).

73. Ayata, C. et al. Laser speckle flowmetry for the study of cerebrovascular physiology in normal and ischemic mouse cortex. J. Cereb. Blood Flow Metab. 24, 744–755 (2004).

74. Aydin, A. K. et al. Transfer functions linking neural calcium to single voxel functional ultrasound signal. Nat. Commun. (2020) doi:10.1038/s41467-020-16774-9.

75. Sans-Dublanc, A. et al. Optogenetic fUSI for brain-wide mapping of neural activity mediating collicular-dependent behaviors. Neuron (2021) doi:10.1016/j.neuron.2021.04.008.

76. Nunez-Elizalde, A. O. et al. Neural basis of functional ultrasound signals. bioRxiv (2021) doi:10.1101/2021.03.31.437915.

77. Brunner, C. et al. Mapping the dynamics of brain perfusion using functional ultrasound in a rat model of transient middle cerebral artery occlusion. J. Cereb. Blood Flow Metab. 37, 263–276 (2017).

78. Brunner, C., Macé, E., Montaldo, G. & Urban, A. Quantitative hemodynamic measurements in cortical vessels using functional ultrasound imaging. Front. Neurosci. 0, p(2022).

79. Rabut, C. et al. 4D functional ultrasound imaging of whole-brain activity in rodents. Nature methods 16, 994–997 (2019).

80. Fisher, M. et al. Update of the stroke therapy academic industry roundtable preclinical recommendations. Stroke 40, 2244–2250 (2009).

