## Supplementary material for "Functional ultrasound imaging of stroke in awake rats": Supp_Figures

Supplementary Figure 1

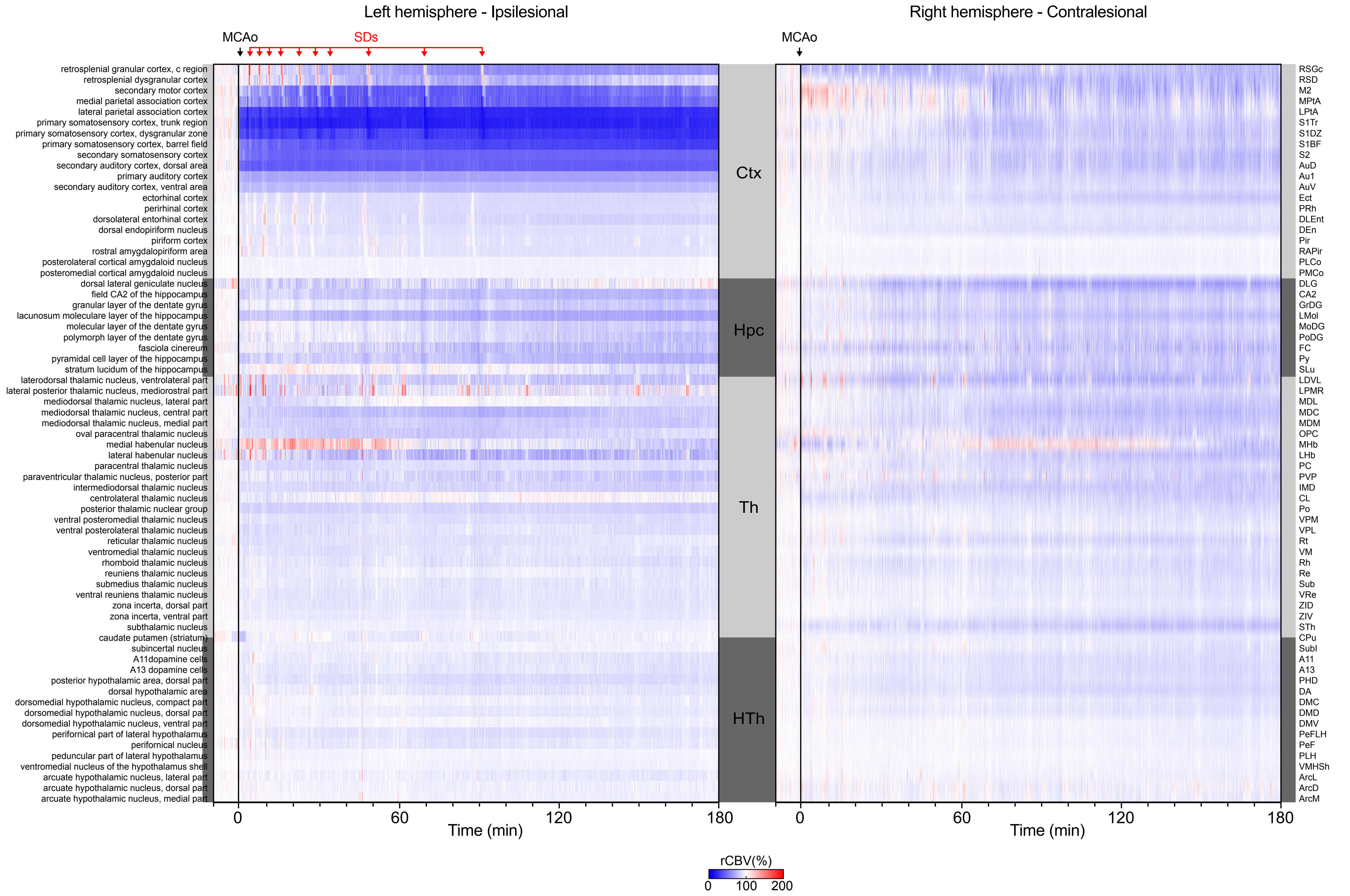

### Supplementary Figure 2

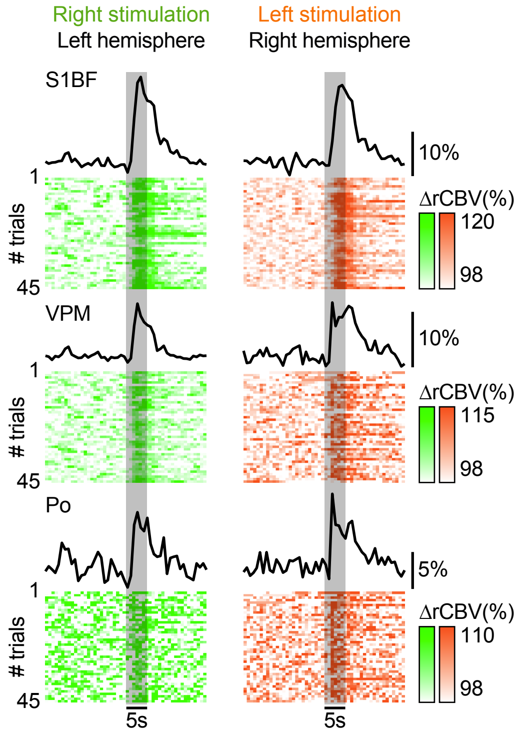

Supplementary Figure 3

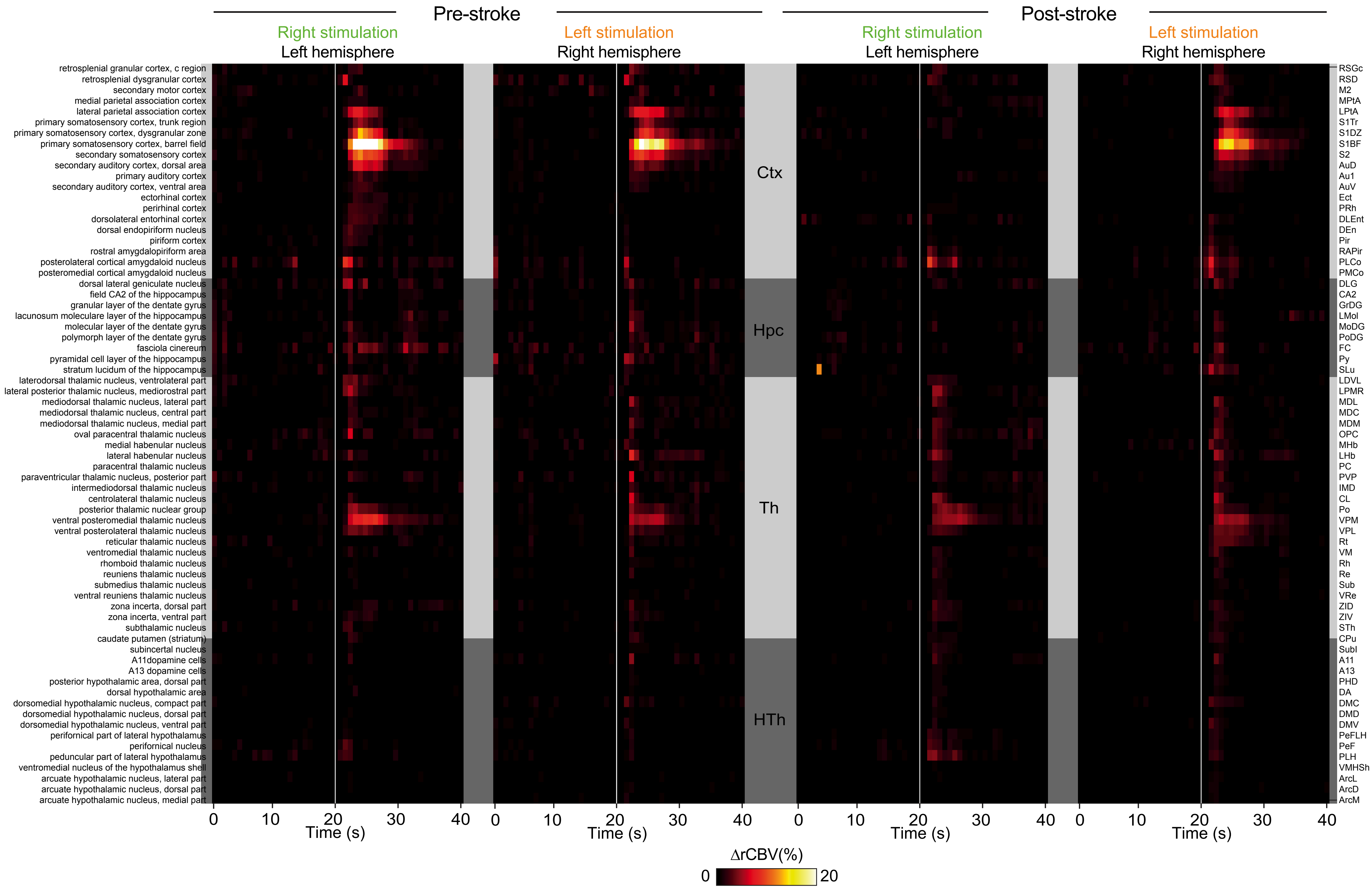

Supplementary Figure 4

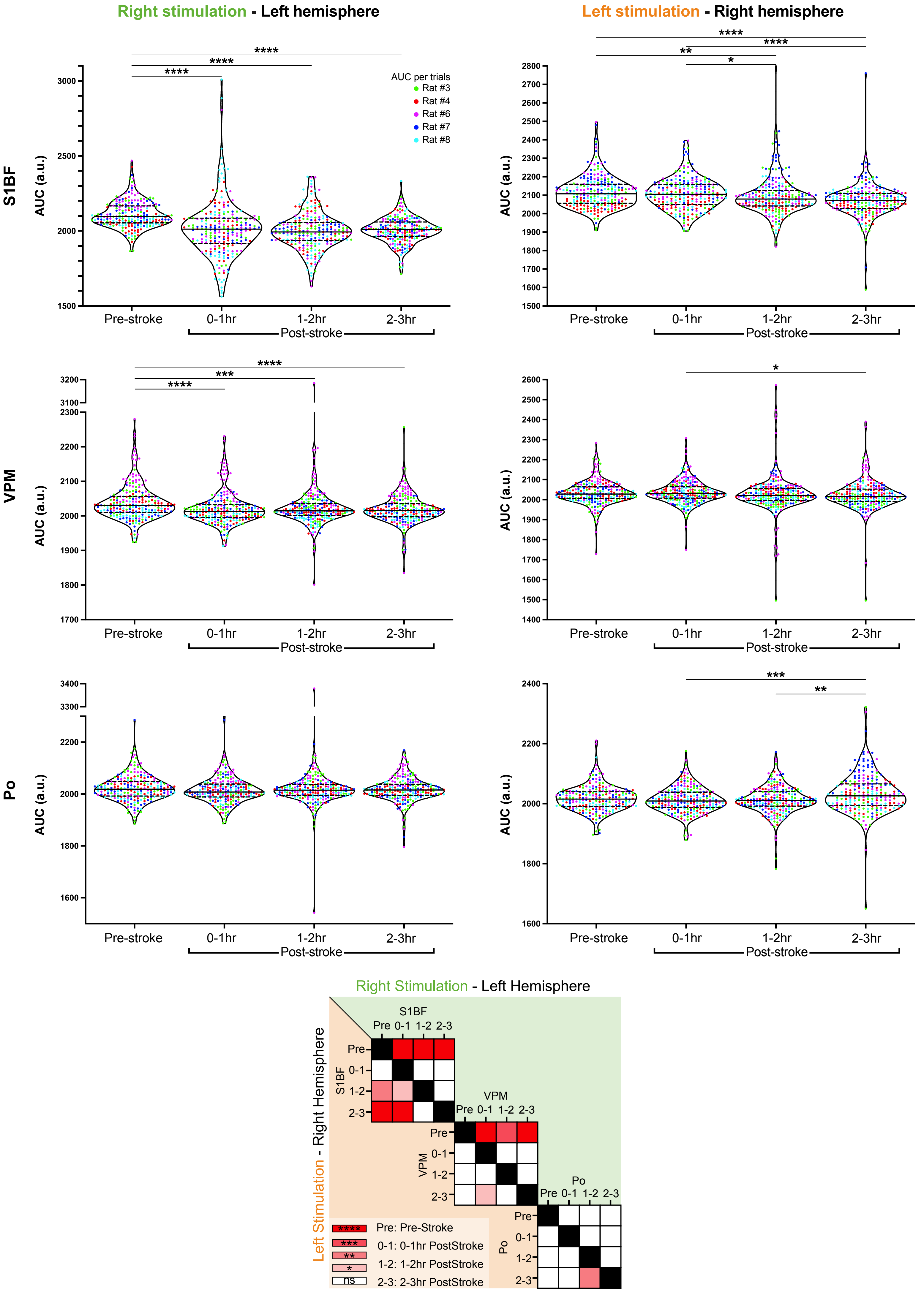

Supplementary Figure 5

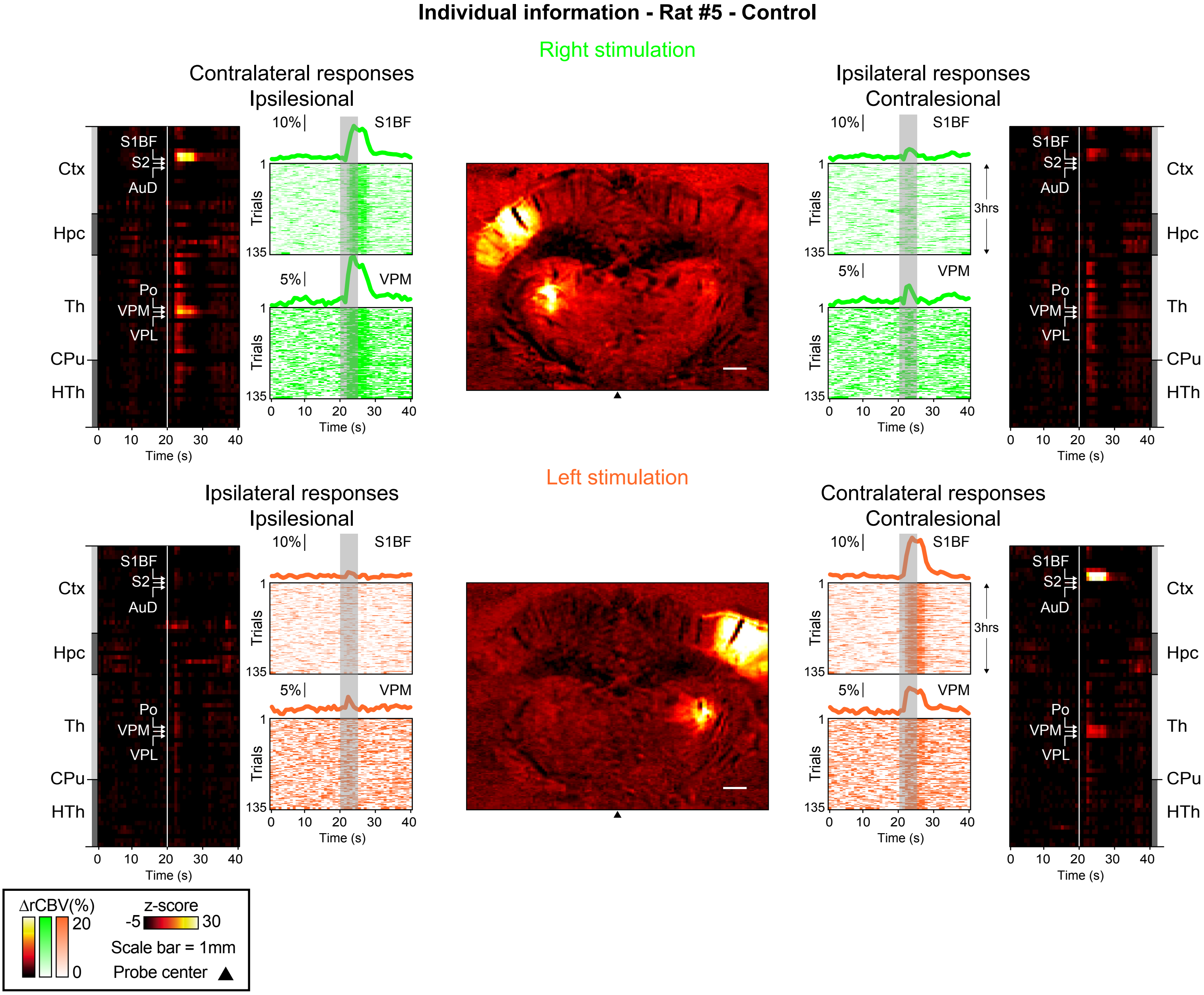

### Individual information - Rat #9 - Control

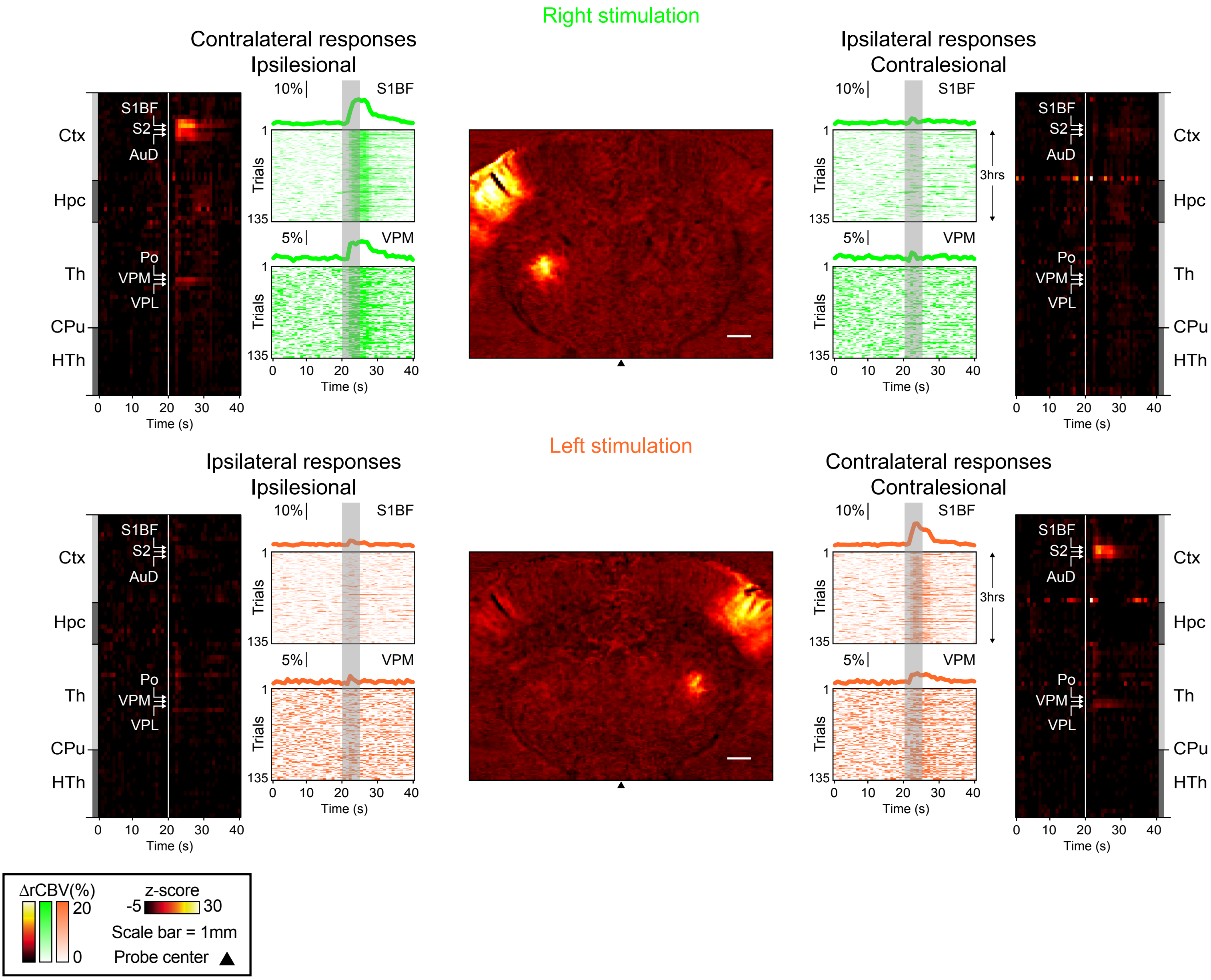

### Individual information - Rat #8 - Control

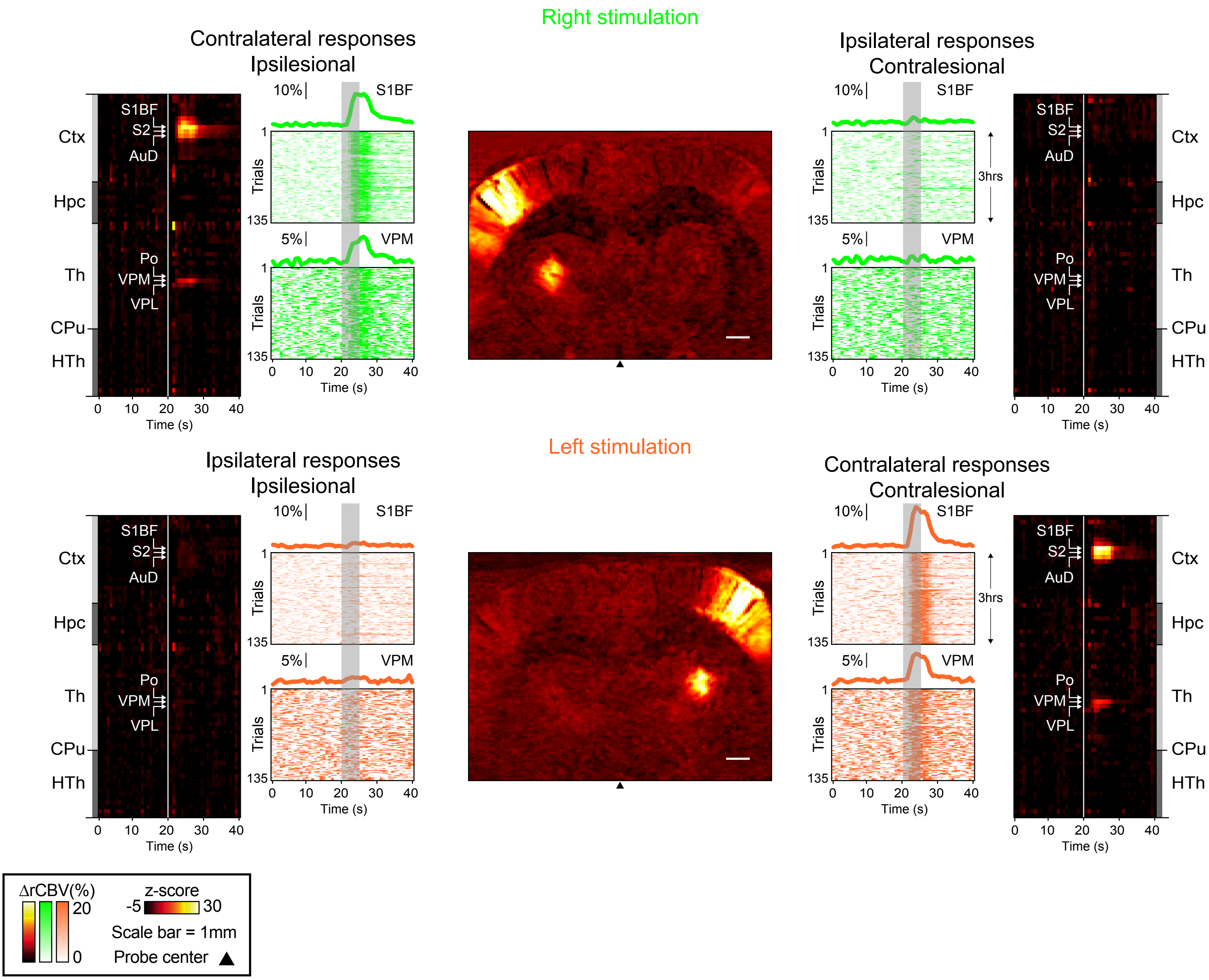

### Individual information - Rat #8 - Pre and Post-Stroke

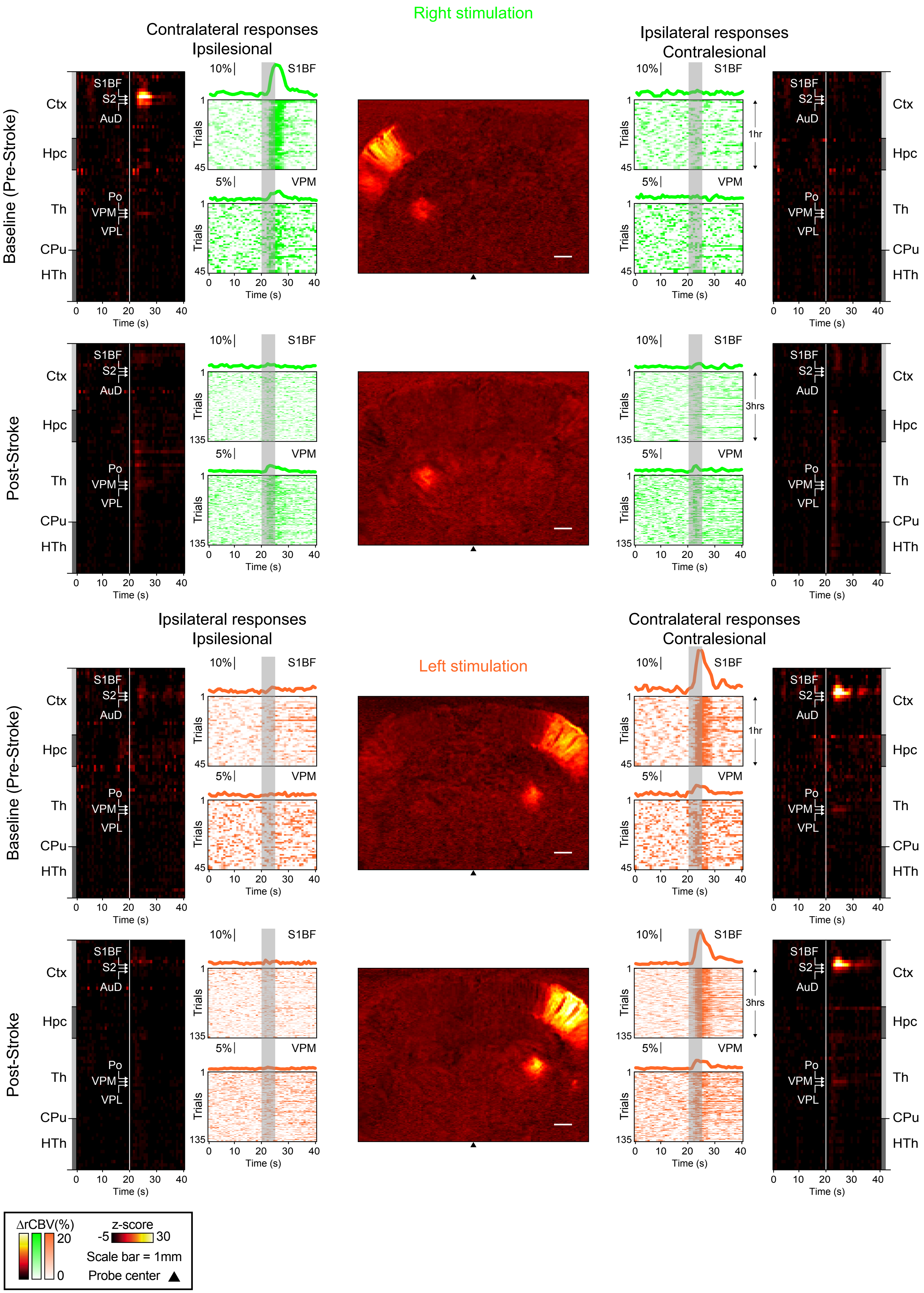

### Individual information - Rat#8 - Post-Stroke 5d

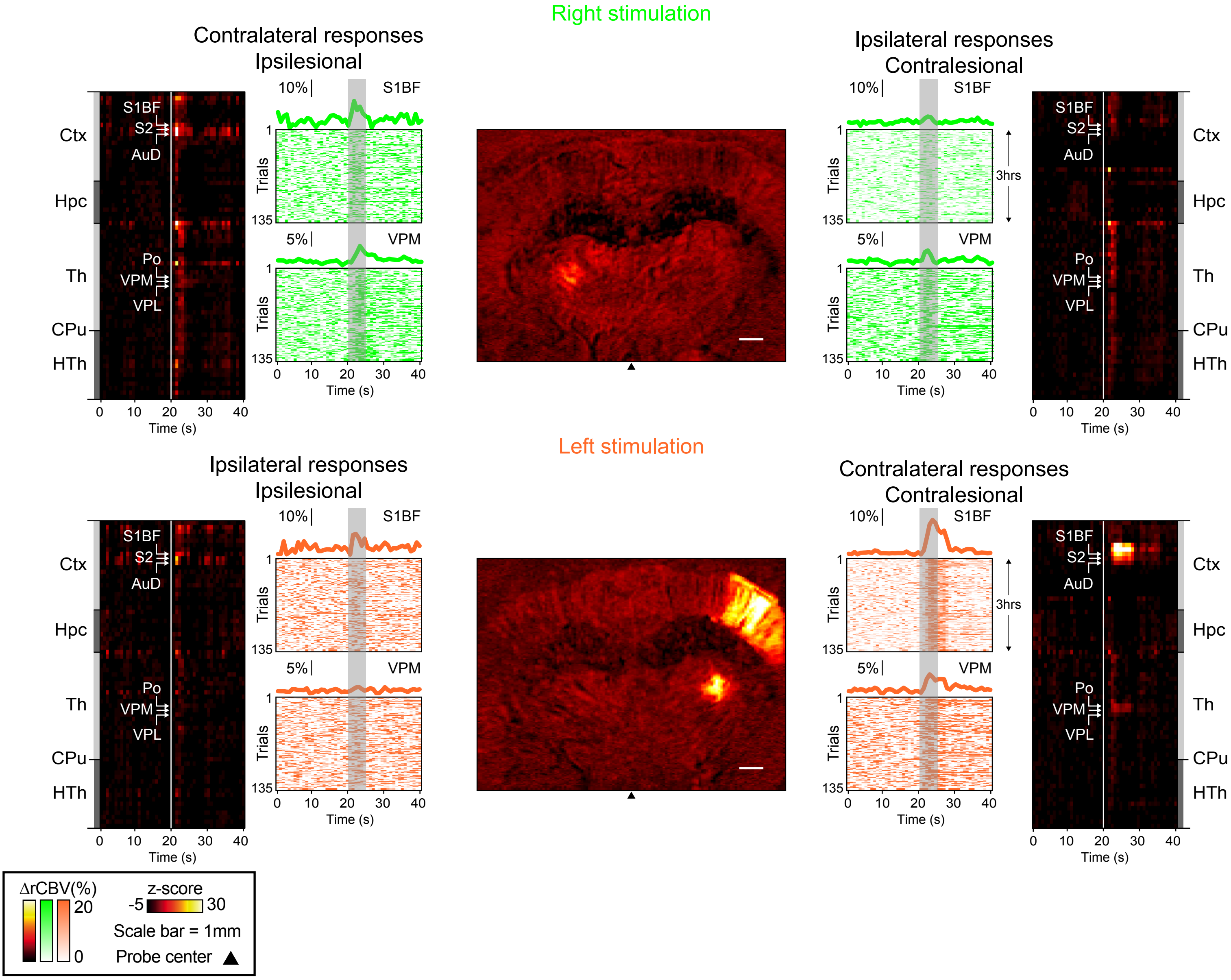

### Individual information - Rat #3 - Pre and PostStroke

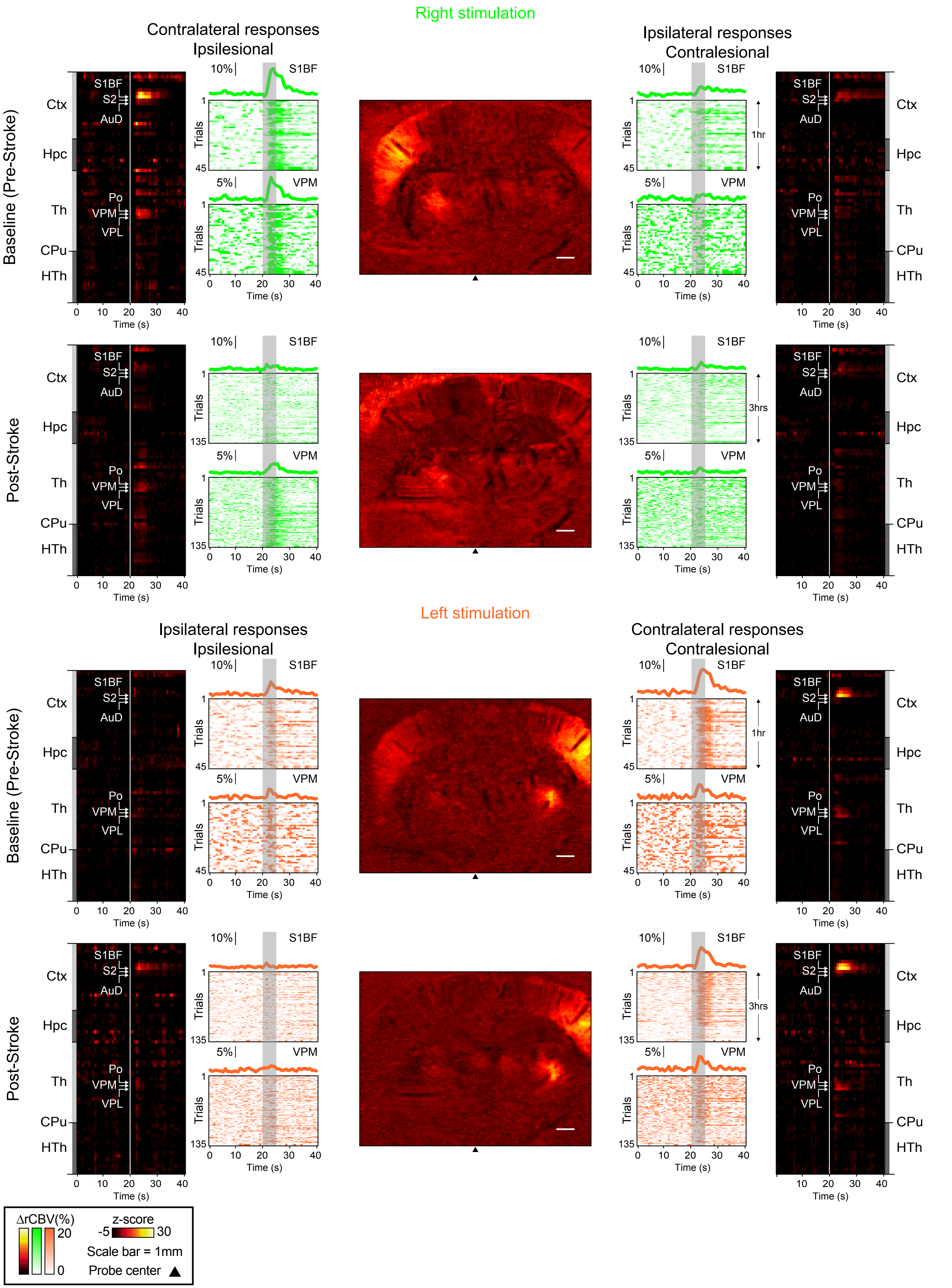

#### Individual information - Rat #4 - Pre and PostStroke

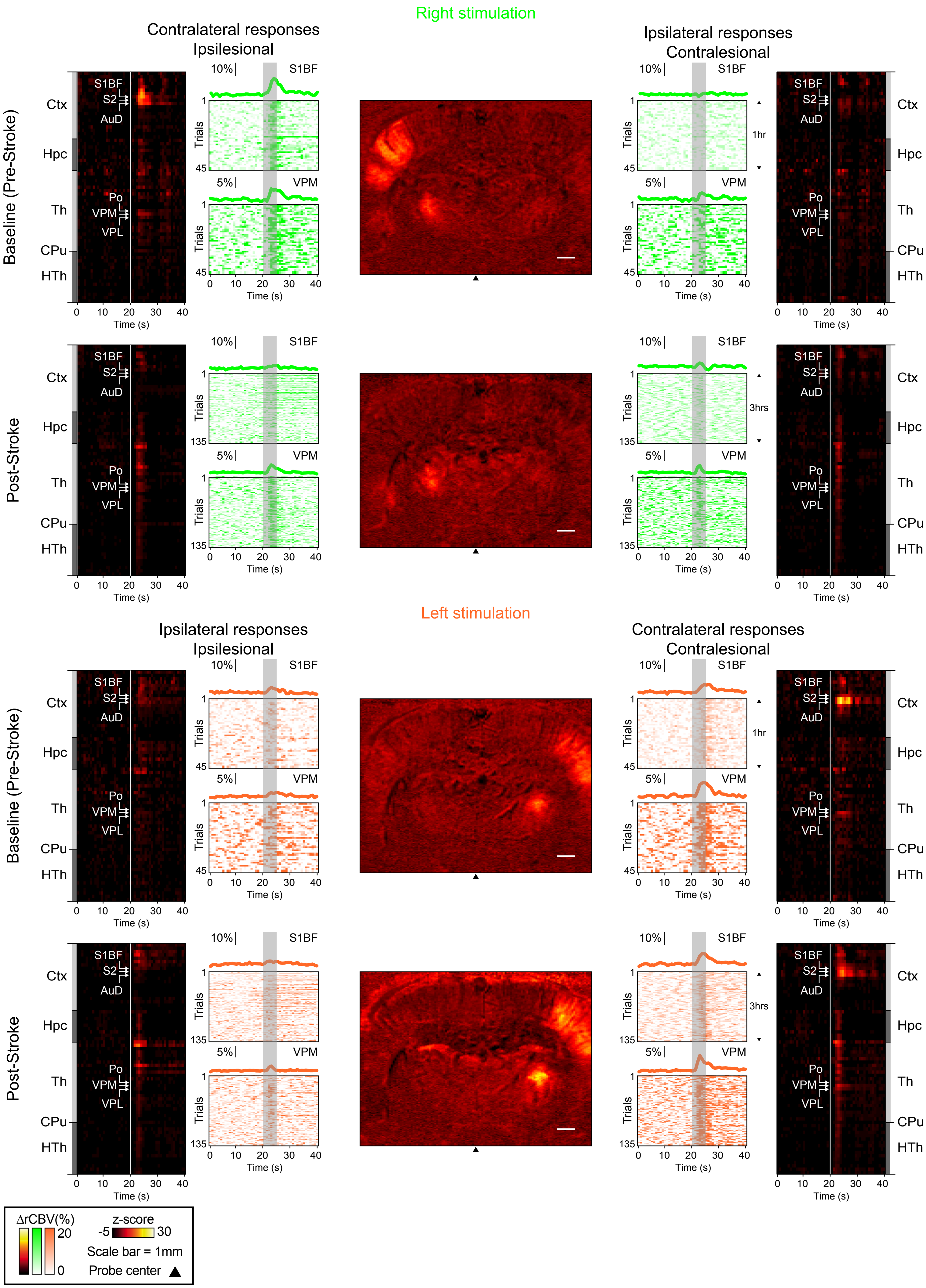

**Individual information - Rat #6 - Pre and PostStroke**

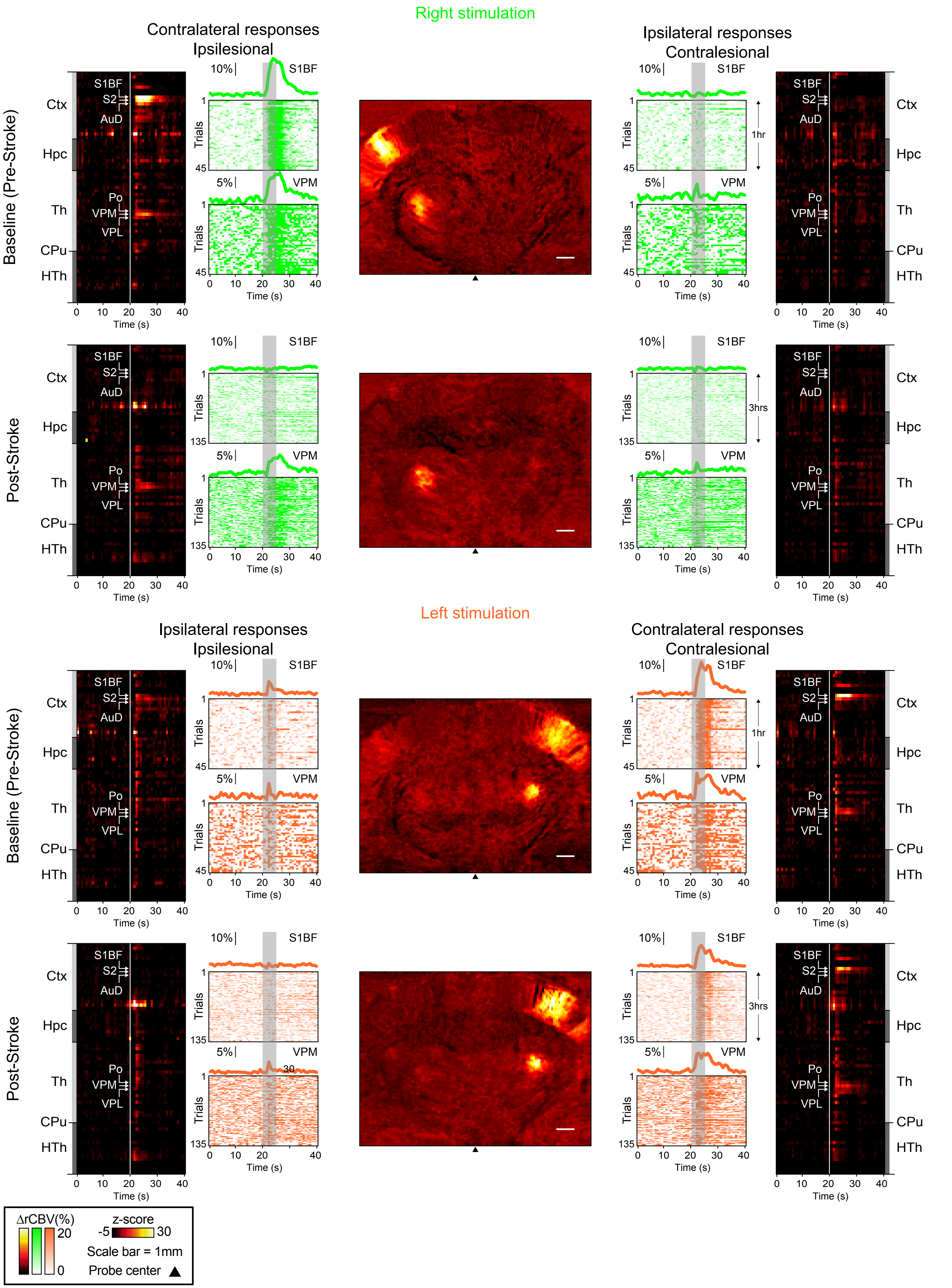

### Individual information - Rat #6 - Post-Stroke 5d

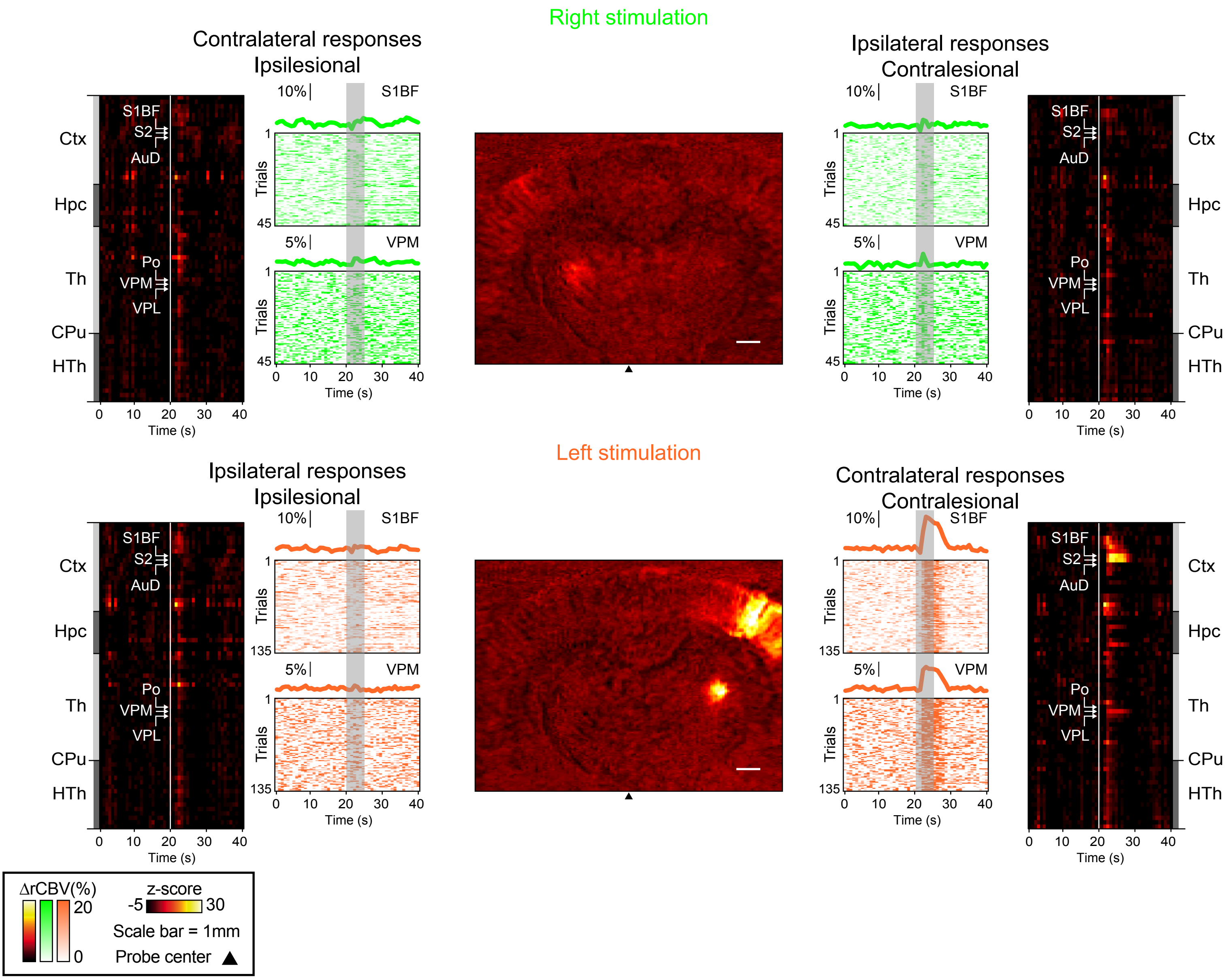

#### Individual information - Rat #7 - Pre and Post-Stroke

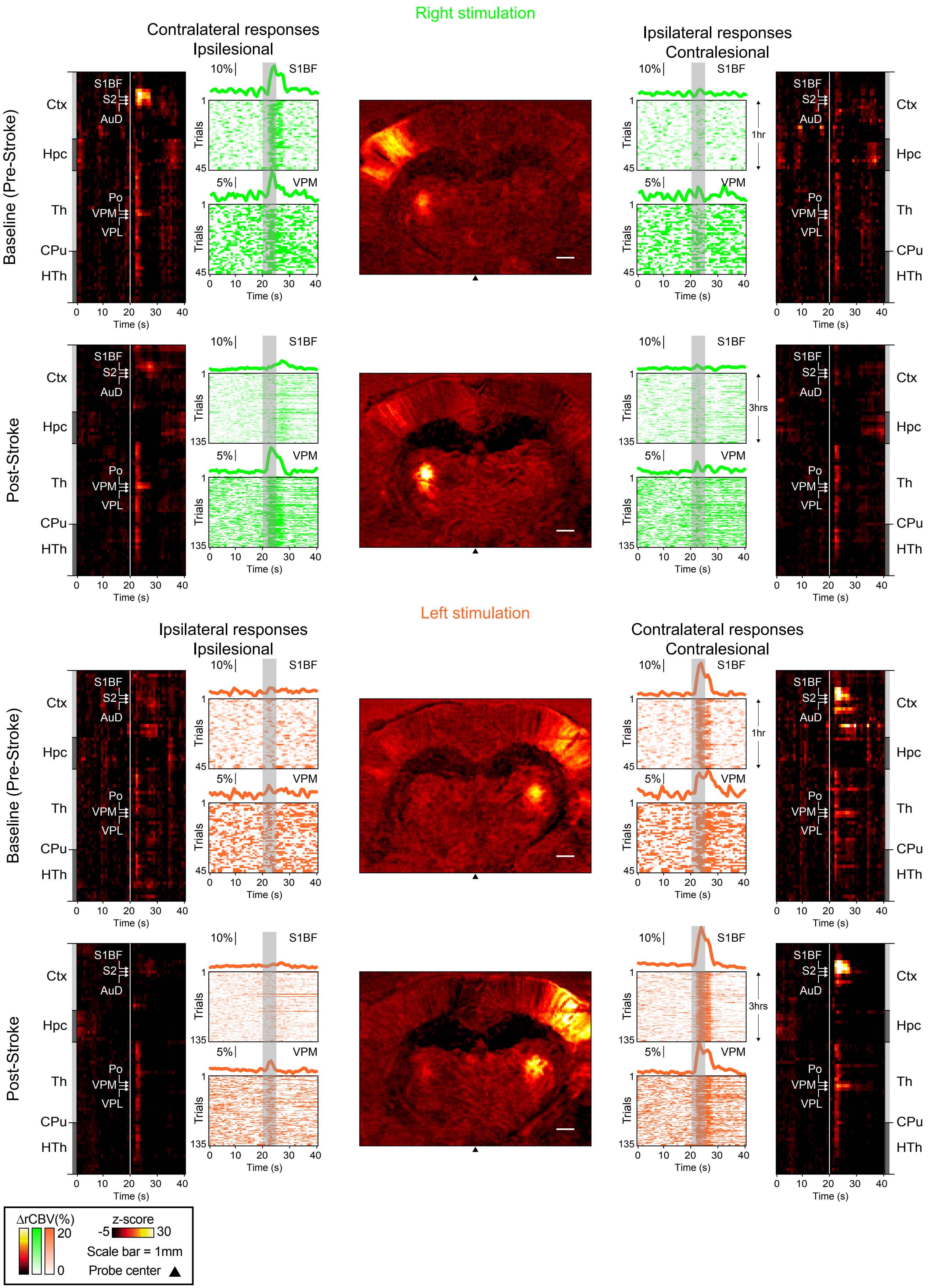
